# Impaired iron recycling from erythrocytes is an early hallmark of aging

**DOI:** 10.1101/2022.01.16.476518

**Authors:** Patryk Slusarczyk, Pratik Kumar Mandal, Gabriela Zurawska, Marta Niklewicz, Komal Kumari Chouhan, Matylda Macias, Aleksandra Szybinska, Magdalena Cybulska, Olga Krawczyk, Sylwia Herman, Michal Mikula, Remigiusz Serwa, Malgorzata Lenartowicz, Wojciech Pokrzywa, Katarzyna Mleczko-Sanecka

## Abstract

Aging affects iron homeostasis, as evidenced by tissue iron loading and toxicity and common anemia in the elderly. Iron needs in mammals are met primarily by iron-recycling from senescent red blood cells (RBCs), a task chiefly accomplished by splenic red pulp macrophages (RPMs) via erythrophagocytosis. Given that RPMs continuously process iron, their cellular functions might be susceptible to age-dependent decline, a condition that has been largely unexplored to date. Here, we found that 10-11-months-old female mice exhibit iron loading, diminished lysosomal activity, and decreased erythrophagocytosis rate in RPMs. These impairments lead to the retention of senescent hemolytic RBCs in the spleen, and the formation of undegradable iron- and heme-rich extracellular protein aggregates, likely derived from ferroptotic RPMs. We further found that feeding mice an iron-reduced diet alleviates iron accumulation in RPMs, enhances their ability to clear erythrocytes, and limits ferroptosis. Consequently, this diet ameliorates hemolysis of splenic RBCs and the formation of iron-rich aggregates, increasing serum iron availability in aging mice. Using RPM-like cells, we show that the diminished iron-recycling capacity of RPMs is underlain by iron accumulation and reduced expression of heme-catabolizing enzyme heme oxygenase 1 (HO-1). Taken together, we identified RPM collapse as an early hallmark of aging and demonstrated that dietary iron reduction improves iron turnover efficacy.

## Introduction

Sufficient iron supplies are critical for cellular metabolic functions and red blood cells (RBCs) production (Cronin et al., 2019; Muckenthaler et al., 2017). As a key constituent of heme - a prosthetic group of hemoglobin, iron confers the unique capability of RBCs to shuffle oxygen and carbon dioxide between the lungs and tissues (Slusarczyk and Mleczko-Sanecka, 2021). At the systemic level, 80% of circulating iron is utilized for hemoglobin synthesis during the daily generation of approximately 200 billion RBCs (Muckenthaler et al., 2017). Hence, in mammals, hemoglobin accounts for the largest pool of heme and iron in the body (Hamza and Dailey, 2012). The oxygen-carrying capacity of RBCs renders them sensitive to the progressive build-up of reactive oxygen species (ROS) that drive natural RBCs senescence (Bratosin et al., 1998). Due to the prooxidative properties of free heme and iron, the exceedingly high hemoglobin content in senescent RBCs constitutes a threat to other cells in the event of RBC breakdown. To reduce the risk of RBCs disintegrating in the blood vessels and because mammals evolved under limited dietary iron availability, 90% of the body iron needs are met by internal iron recycling from aged RBCs (Ganz, 2012). This task is accomplished predominantly by red pulp macrophages (RPMs) of the spleen, cells that recognize and engulf defective RBCs in the process called erythrophagocytosis (EP) (Bian et al., 2016; Youssef et al., 2018).

The loss of elasticity is a key feature of naturally aged RBCs (Bratosin et al., 1998; Higgins, 2015; Lutz, 2012). One of the underlying mechanisms is the oxidation and denaturation of hemoglobin, as well as the formation of lipid peroxides (Arashiki et al., 2013; Ganz, 2012). Once RBCs become too rigid to pass through slits within the red pulp’s unique venous system, they are retained within the spleen cords (Mebius and Kraal, 2005). Recognition of “trapped” RBCs by RPMs involves additional signals proposed to act additively (Gottlieb et al., 2012; Slusarczyk and Mleczko-Sanecka, 2021). Upon engulfment by RPMs, RBCs are degraded in phagolysosomes, globins are hydrolyzed to amino acids, and heme is released (Klei et al., 2017). Recent findings suggest that in addition to phagocytosis, part of the RBC-derived iron is recovered via hemolysis of RBCs within the splenic microenvironment and subsequent endocytic hemoglobin uptake (Klei et al., 2020). In RPMs, heme is transported from phagolysosomes or late endosomes to the cytoplasm by HRG1 (Pek et al., 2019) and subsequently catabolized by heme oxygenase 1 (HO-1; encoded by *Hmox-1*) to carbon monoxide (CO), biliverdin, and ferrous iron that feeds the labile iron pool (LIP) (Kovtunovych et al., 2010). The nanocage-like protein ferritin sequesters iron from the LIP. Iron efflux from RPMs occurs via ferroportin (FPN) to replenish the transferrin-bound iron pool in the plasma (Muckenthaler et al., 2017; Zhang et al., 2011). The process of iron release from RPMs is tightly regulated by hepcidin, a small liver-derived hormone that mediates FPN degradation and/or occlusion, hence preventing iron release from the RPMs iron reservoir (Aschemeyer et al., 2018; Nemeth et al., 2004). Despite the growing body of knowledge, it is still insufficiently explored how iron balance in RPMs is affected by different pathophysiological conditions and how it may influence their functions.

Similar to erythropoiesis, the rate of RBCs sequestration is very high, reaching 2-3 million per second in humans (Higgins, 2015), and mainly considered constitutive under physiological conditions. Some reports showed that the intensity of EP can be enhanced by proinflammatory conditions and pathogen components (Bennett et al., 2019; Bian et al., 2016; Delaby et al., 2012). Recent work employing genetic mouse models showed that altered calcium signaling in RPMs due to overactivation of the PIEZO1 mechanoreceptor (Ma et al., 2021) or deficiency of the IL-33 receptor IL1RL1 (Lu et al., 2020), induces or represses EP rate, respectively. However, it is largely unknown if the RBCs uptake and degradation within the phagolysosomes are regulated by intrinsic or systemic iron status *per se*.

Iron dyshomeostasis hallmarks physiological aging. This is exemplified by progressive iron accumulation and iron-dependent oxidative damage in aging organs, such as the brain, liver, spleen, kidney, skeletal muscles, or heart (Arruda et al., 2013; Cook and Yu, 1998; Sukumaran et al., 2017; Xu et al., 2008). At the same time, plasma iron deficiency is frequent in aged individuals and is a leading cause of a condition referred to as anemia in the elderly (Girelli et al., 2018). RPMs are derived from embryonic progenitor cells (Yona et al., 2013), exhibit limited self-renewal capacity (Hashimoto et al., 2013), and are only partially replenished by blood monocytes during aging (Liu et al., 2019). Hence, these specialized cells continuously perform senescent RBCs clearance during their lifespan, thus processing and providing most of the iron the organism utilizes. We hypothesized that this might render RPMs particularly susceptible to age-related iron-triggered functional deterioration, affecting body iron indices and RBCs homeostasis. Here, we found that aging impairs the capacity of RPMs to engulf and lyse RBCs and promotes their loss, likely via ferroptosis. We propose that these defects affect local RBCs’ balance in the spleen, augmenting the degree of their lysis, and promote the formation of iron- and heme-rich extracellular protein aggregates. We decipher new mechanisms of EP regulation by an interplay between intracellular iron levels, HO-1 activity, and proteotoxic endoplasmic reticulum (ER) stress. Finally, we provide evidence that reducing dietary iron during aging “rejuvenates” RPMs function and improves iron homeostasis.

## Results

### RPMs of aged mice show increased labile iron levels, oxidative stress and diminished iron-recycling functions

To investigate iron recycling capacity during aging, we used 10-11-month-old mice fed a diet with standard iron content (200 ppm). We used female mice as they show a higher iron burden in the spleen with age progression (Altamura et al., 2014). This age in mice corresponds to approximately 40-45 years 31 in humans when senescent changes begin to occur (Fox, 2006). As expected, aged mice show tissue iron loading, with the spleen being most affected, followed by the liver and other organs such as muscles and the heart (Fig. S1A-D). Aging mice exhibit decreased blood hemoglobin values, as previously shown (Peters et al., 2008), and a drop in transferrin saturation, a feature that thus far was observed mostly in elderly humans (Fig. S1E and F) (Girelli et al., 2018). Next, we aimed to verify if aging affects the iron status and essential cellular functions of RPMs. Ferritin is an iron-storing heteropolymer composed of H and L subunits. Due to its ferroxidase activity, H ferritin is a key endogenous protective protein that converts redox-active ferrous iron to inert ferric iron retained inside the ferritin shells (Mleczko-Sanecka and Silvestri, 2021). Using intracellular staining and flow cytometric analyses, we uncovered that RPMs [gated as CD11b-dim, F4/80-high, TREML4-high (Haldar et al., 2014); Fig. S2] in aged mice exhibit a significant deficiency in H ferritin levels, with unchanged L ferritin protein expression (Fig. S1G and H). Consistently, using the fluorescent probe FerroOrange that interacts explicitly with ferrous iron, we detected a significant increase in LIP in aged RPMs (Fig. S1I) accompanied by marked oxidative stress (detected using CellROX probe; Fig. S1J). We next tested if this increase in labile iron in RPMs would impact the phagocytic activity. Hence, we incubated splenic single-cell suspension with temperature-stressed RBCs fluorescently labeled with PKH67 dye (Klei et al., 2020; Theurl et al., 2016). In parallel, we employed another standard cargo for phagocytosis, zymosan, a yeast cell wall component. Using this *ex vivo* approach, we detected a significant drop in RBCs clearance rate in aged compared to young RPMs (Fig. S1K). Notably, their capacity for the engulfment of zymosan remained unchanged (Fig. S1L), thus suggesting a specific defect of EP in aged RPMs. Using a fluorescent probe, we also observed decreased lysosomal activity in RPMs isolated from 10-11-months old mice compared to those derived from young control animals (Fig. S1M). Interestingly, peritoneal macrophages of aged mice did not show altered ROS levels or diminished functions of lysosomes and mitochondria (Fig. S3), suggesting that these age-related changes affect RPMs earlier than other macrophage populations. Altogether, these insights suggest that during aging, RPMs increase LIP and exhibit a reduced capacity for the RBCs engulfment and lysosomal degradation of erythrocyte components.

### Iron-reduced diet normalizes body iron parameters during aging, diminishes iron retention in RPMs, and rescues age-related decline in their iron-recycling capacity

We next set out to explore whether an age-related redistribution of iron from plasma to organs, particularly the spleen, could be prevented. Caloric restriction was previously shown to effectively reduce tissue iron content in older rats in organs such as muscle, kidney, liver, and brain (Cook and Yu, 1998; Xu et al., 2008). To target iron dyshomeostasis in aging mice, we applied an alternative approach by reducing dietary iron from the fifth week of life to a level (25 ppm) that was reported as sufficient to maintain erythropoiesis (Sorbie and Valberg, 1974). This approach reverted the degree of iron deposition in the spleen and liver (Fig. 1A and B) and partially alleviated the decreased transferrin saturation characteristic of aged mice fed a standard diet (Fig. 1C). Notably, the mild drop in blood hemoglobin was not rescued by the iron-reduced (IR) diet (Fig. S4A). Consistently, extramedullary erythropoiesis in the spleen and increased plasma erythropoietin (EPO) levels in aged mice were not affected by dietary iron content (Fig. S4B and C). These observations imply that partial alleviation of plasma iron availability in IR mice is insufficient to correct mild aging-triggered anemia and/or other hematopoiesis-related mechanisms may contribute to this phenotype. Next, we observed that hepcidin mRNA levels in the liver reflected, as expected, the degree of hepatic iron deposition in mice that were aged on the standard and IR diets (Fig. 1D). The alterations in hepcidin expression levels were reflected by the changes in FPN levels on the surface of RPMs, as indicated by flow cytometry analyses using an antibody that recognizes the extracellular loop of murine FPN (Fig. 1E) (Sangkhae et al., 2019). Hence, we further characterized RPMs’ iron status. We found that despite decreased ferritin H levels in aged mice, regardless of the dietary regimen (Fig. 1F), accumulation of both labile iron (measured using FerroOrange and flow cytometry; Fig. 1G) and total iron (measured by colorimetric assay in magnetically-sorted RPMs, Fig. 1H) is completely reversed in RPMs in aged mice fed an IR diet as compared to a standard diet. Likewise, we observed alleviation of oxidative stress in RPMs, using a pan-ROS fluorescent probe CellROX (Fig. 1I).

**Figure 1.**
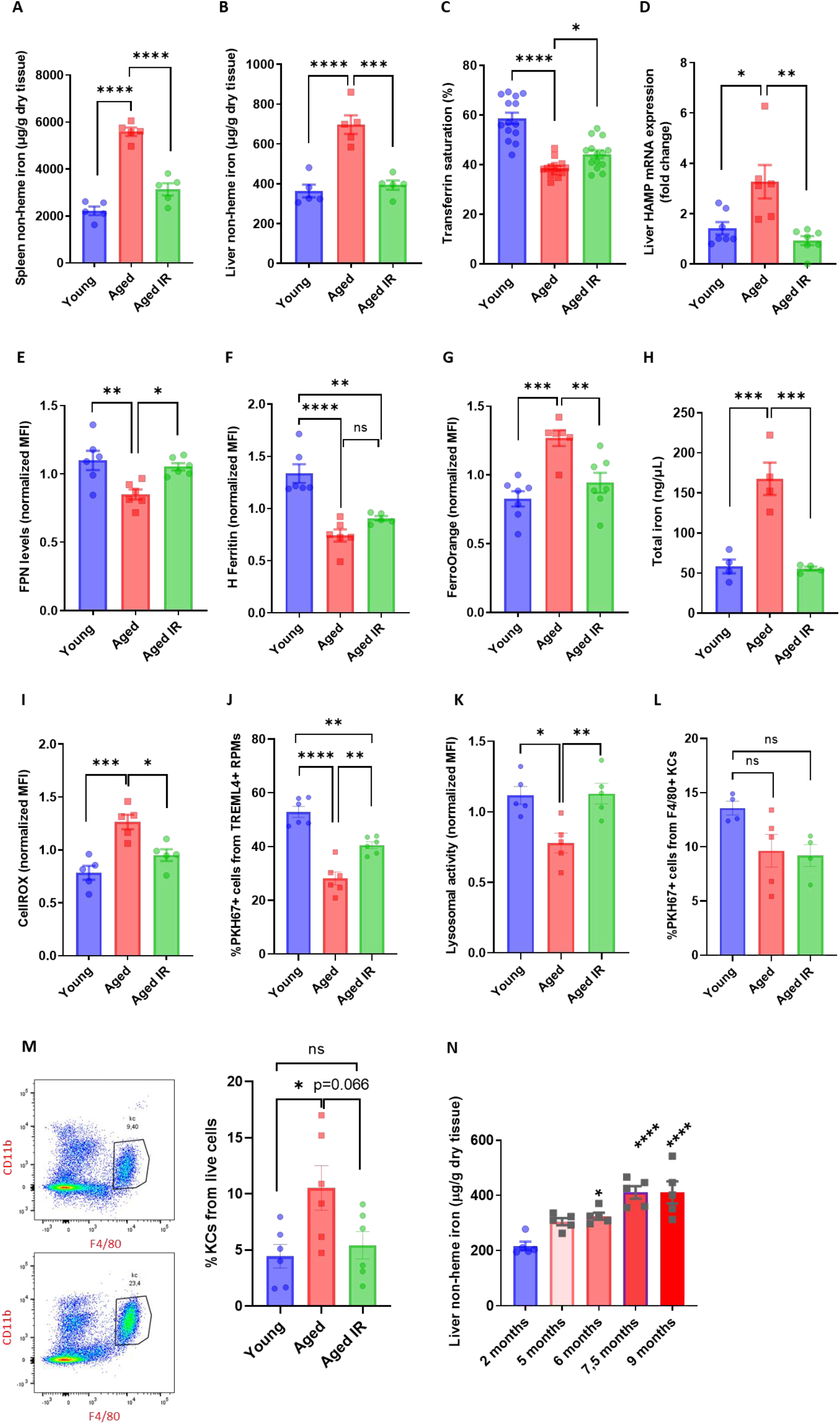
Iron-reduced diet normalizes body iron parameters during aging, diminishes iron retention in RPMs, and rescues age-related decline in their iron-recycling capacity. (A) Splenic and (B) hepatic non-heme iron content was determined in young, aged, and aged IR mice. (C) Plasma transferrin saturation was determined in young, aged, and aged IR mice. (D) Relative mRNA expression of hepcidin (Hamp) in the liver was determined by qPCR. (E) Cell surface ferroportin expression in young, aged, and aged IR RPMs was assessed by flow cytometry. (F) H-Ferritin protein levels in young, aged, and aged IR RPMs were quantified by flow cytometric analysis. (G) Cytosolic ferrous iron (Fe2+) levels in young, aged, and aged IR RPMs were measured using FerroOrange and flow cytometry. (H) The total intracellular iron content in young, aged, and aged IR magnetically-sorter RPMs was assessed using a commercial assay and normalized to cell number. (I) The cytosolic ROS levels in young, aged, and aged IR RPMs were assessed by determining CellROX Deep Red fluorescence intensity with flow cytometry. (J) RBC clearance capacity in young, aged, and aged IR RPMs was determined by measuring the percentage of RPMs that phagocytosed transfused PKH67-labeled temperature-stressed RBCs. (K) The lysosomal activity of young, aged, and aged IR RPMs was determined using a dedicated fluorescent probe and flow cytometry. (L) The phagocytosis rate of transfused PKH67-labeled temperature-stressed RBCs by Kupffer cells (KCs) in young, aged, and aged IR mice was measured with flow cytometry analysis. (M) Representative flow cytometry plots of KCs and percentages of KCs out of total live cells in livers of young, aged, and aged IR mice. (N) Liver non-heme iron content was determined in mice at the indicated age. Each dot represents one mouse. Data are represented as mean ± SEM. Statistical significance among the three or more groups was determined by One-Way ANOVA test with Tukey’s Multiple Comparison test. *p<=0.05, **p<=0.01, ***p<=0.001 and ****p<=0.0001

The major function of RPMs is the clearance of aged erythrocytes. We thus explored if their altered iron status in aged mice reflects their capacity to engulf and degrade RBCs. To this end, we first performed *in vivo* erythrophagocytosis assay via transfusion of PKH67-stained temperature-stressed erythrocytes (Lu et al., 2020; Theurl et al., 2016). We identified a significant reduction in the capacity of RPMs to sequester RBCs in aged mice, an impairment that was partially rescued by feeding mice an IR diet (Fig. 1J). Similarly to the EP rate, we found that RPMs isolated from aged IR mice restored the lysosomal activity to the levels observed in young mice (Fig. 1K). These data imply that the diminished capacity of RPMs to sequester and lyse RBCs during aging can be ameliorated by limiting dietary iron content. Since earlier studies demonstrated that under conditions of RPMs impairment, iron-recycling functions are significantly supported by liver macrophages (Lu et al., 2020; Theurl et al., 2016), we also measured their EP rate during aging. Using the transfusion of PKH67-stained RBCs (Akilesh et al., 2019) we found no increase in Kupffer cells (KCs) EP capacity in mice fed IR (Fig. 1L). However, consistently with the *de novo* recruitment of iron-recycling myeloid cells to the liver upon erythrocytic stress (Theurl et al., 2016), we detected a significant expansion of the KC-like population in aged mice but not in the aged IR group (Fig. 1M), suggesting that by this means, the liver macrophage may partially compensate for the reduced EP activity in RPMs. In support of the model that “on-demand” hepatic iron-recycling macrophages support iron recycling during aging, and in agreement with the previous findings that iron from engulfed RBCs is stored in hepatocytes (Theurl et al., 2016), we observed progressive liver iron accumulation with age (Fig. 1N). In sum, we demonstrate that dietary iron restriction normalizes both systemic and RPMs iron levels, thus preventing the onset of cellular oxidative stress in RPMs of aged mice and rescuing their age-related decline in iron-recycling capacity.

### Aging triggers retention of senescent RBCs, increased hemolysis, and the formation of non-degradable iron-rich aggregates in the spleen

Having established that dietary iron content in aging modulates RPMs EP capacity (Fig. 1J), we examined parameters related to RBCs fitness. First, we performed an RBCs lifespan assay and found no differences in the rate of RBCs removal from the circulation between young and aged mice (Fig. 2A). However, defective RBCs are first filtered in the spleen due to the loss of their elasticity, and this step is a prerequisite for their removal via EP (Slusarczyk and Mleczko-Sanecka, 2021). In agreement with this model, RBCs retained in the spleen of young mice show ROS build-up, a marker of their senescence (Bratosin et al., 1998), compared to the circulating RBCs (Fig. 2B and S5). We thus hypothesized that reduced phagocytic capacity of RPMs in aging might impact local rather than systemic RBCs homeostasis. Consistently, splenic RBCs isolated from older mice show more pronounced ROS levels than RBCs derived from young mice (Fig. 2C). In line with the partial mitigation of the defective EP capacity (Fig. 1J), we found that mice fed an IR diet exhibited erythrocytic oxidative stress parameters as the young mice (Fig. 2D). Since ROS can accumulate in RBCs with deficient FPN (Zhang et al., 2018), we examined FPN levels of splenic RBCs. We found no reduction but a significant FPN increase in aged mice, implying that increased erythrocytic ROS are not caused by alterations in FPN levels (Fig. 2E). Furthermore, in agreement with the notion that a part of senescent splenic RBCs undergoes hemolysis (Klei et al., 2020), we detected significantly higher levels of extracellular heme in the aged spleens, likely reflecting a higher burden of defective RBCs that escape EP (Fig. 2F). Interestingly, we found that the levels of this free heme in aging spleens are partially rescued by the IR diet. In conclusion, our data suggest that reduced EP rate during aging may promote the splenic retention of senescent RBCs prone to undergo local lysis and that these physiological changes may be partially rescued by maintaining mice on an IR diet.

**Figure 2.**
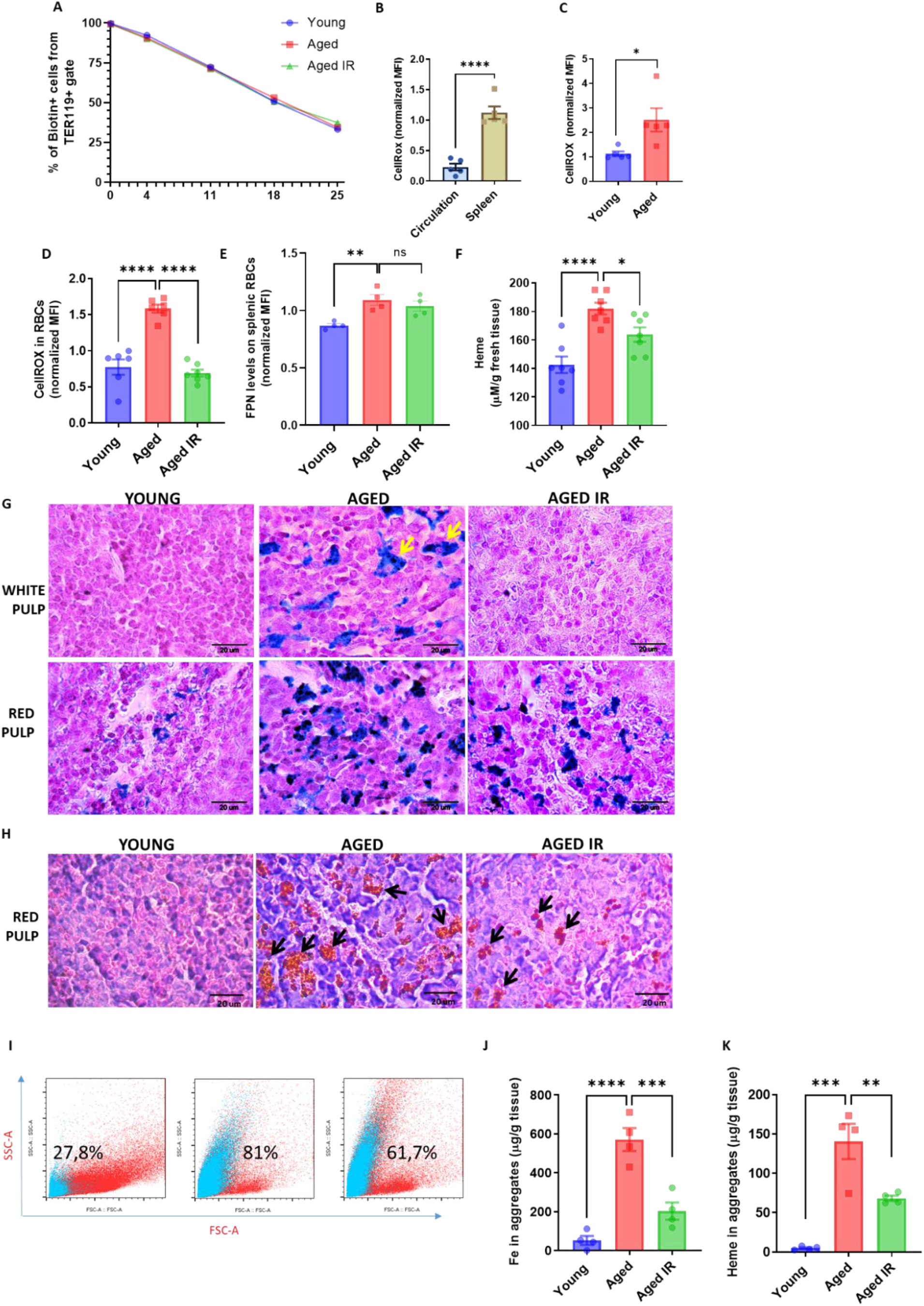
Aging triggers retention of senescent RBCs, increased hemolysis, and the formation of non-degradable iron-rich aggregates in the spleen. (A) The RBC biotinylation assay was performed on circulating RBCs from young, aged, and aged IR mice. (B–D) The cytosolic ROS levels in RBCs derived from (B) circulation and the spleen of young mice; (C) the spleen of young and aged mice; (D) the spleen of young, aged, and aged IR mice. Levels of ROS were estimated by determining CellROX Deep Red fluorescence intensity with flow cytometry. (E) Expression of ferroportin on the surface of RBCs obtained from the spleen of young, aged, and aged IR mice assessed by flow cytometry. (F) Extracellular heme levels were measured in the medium obtained after dissociation of spleens from young, aged, and aged IR mice using Heme Assay Kit. (G) Perls’ Prussian Blue staining of the splenic white and red pulp in young, aged, and aged IR mice. Arrows indicate cells with ferric iron deposits in the white pulp. (H) Hematoxylin/eosin staining of the splenic red pulp in young, aged, and aged IR mice. Arrows indicate extracellular brown aggregates. (I) Representative flow cytometry plots of magnetically-sorted splenocytes. In blue are events adverse for dead cell staining and cell surface markers CD45, TER119, CD11b, and F4/80. The percentage of these events in total acquired events is indicated. (J) The total iron content in magnetically-sorted, cell-free aggregates derived from spleens of young, aged, and aged IR RPMs mice was assessed using a commercial assay. (K) The heme content in magnetically-sorted, cell-free aggregates derived from spleens of young, aged, and aged IR RPMs mice was assessed using Heme Assay Kit. Each dot in the graph (A) represents n=8. For the other panels, each dot represents one mouse. Data are represented as mean ± SEM. Welch’s unpaired t-test determined statistical significance between the two groups; for the three groups, One-Way ANOVA with Tukey’s Multiple Comparison test was applied. *p<=0.05, **p<=0.01, ***p<=0.001 and ****p<=0.0001

Previous work showed that genetic deletion of SPI-C, the transcription factor that confers RPMs functional identity (Haldar et al., 2014; Kohyama et al., 2009), leads to the gradual loss of RPMs in young postnatal mice (Okreglicka et al., 2021). The knock-out of *Pparg*, a factor required for neonatal RPM expansion, is manifested by a profound depletion of RPMs (Okreglicka et al., 2021), whereas low numbers of fully differentiated RPMs and their compromised EP capacity hallmark the genetic abrogation of the IL-33-IL1RL1 pathway (Lu et al., 2020). A common denominator for all these distinct mouse models is iron accumulation in the spleen, implying that splenic iron overload may result from diminished RPMs functionality. We thus aimed to explore the identity of splenic iron deposits in more detail. Perls’ Prussian blue staining of spleen sections showed that aged mice on a standard diet exhibit enhanced iron accumulation in the red pulp, a less pronounced phenotype in mice fed an IR diet (Fig. 2G). This primarily reflects RPMs iron status (Fig. 1G and H). We also detected dispersed large iron-loaded cells in the white pulp of aged mice fed a standard diet which were absent in young and iron-reduced mice (Fig. 2G). Intriguingly, eosin/hematoxylin staining visualized deposits in aged spleens that, in contrast to Perls’ staining, were completely absent in young controls (Fig. 2H). They appear as large and extracellular in the histology sections and seem smaller and less abundant in mice fed an IR diet (Fig. 2H). Flow cytometry analyses confirmed that splenic single-cell suspension from aged but not young mice contains superparamagnetic particles, likely rich in iron oxide (Franken et al., 2015), that are characterized by high granularity as indicated by the SSC-A signal. These particles do not show staining indicative of dead cells and fail to express typical splenocyte and erythroid markers (CD45, TER119, CD11b or F4/80) (Fig. 2I). With different granularity and size parameters, such particles appear as well in mice aged on an IR diet, however, to a lesser extent. We established a strategy involving splenocyte separation in the lymphocyte separation medium followed by magnetic sorting, which successfully separates iron-rich cells (chiefly RPMs) from iron-containing extracellular aggregates (Fig. S6). We found that these aggregates contain large amounts of total iron (Fig. 2J) and, interestingly, heme (Fig. 2K), implying their partial RBC origin. Importantly, both iron and heme content is significantly reduced in mice fed an IR diet. In sum, our data show that aging is associated with local RBC dyshomeostasis in the spleen, and the formation of iron- and heme-rich aggregates, and that both these responses can be alleviated by limiting dietary iron content.

### Splenic age–triggered iron deposits are rich in aggregation-prone protein debris that likely emerges from dysfunctional and damaged RPMs

We next sought to explore the origin of the splenic iron aggregates. We imaged spleen sections from young and aged mice with transmission electron microscopy (TEM) to obtain ultrastructural information. We noticed that RPMs in young mice contain only intracellular dark-colored deposits, rich in structures that resemble ferritin (Fig.3 Ai). Their appearance likely mirrors a form of iron deposits that were in the past referred to as hemosiderin, an intracellular clustered insoluble degradation product of ferritin (Harrison and Arosio, 1996; Ward et al., 2000). In older mice, we likewise observed intracellular deposits in morphologically vital cells (Fig. 3Aii). However, in agreement with histological staining, we detected two other classes of aggregates: those that are still granular and enclosed within the membrane, but present in cells that are morphologically damaged (Fig. 3Aiii) and those that are large, amorphic, and located extracellularly (Fig. 3Aiv). Next, we conducted label-free proteomic profiling to determine the composition of magnetically-isolated aging-associated splenic aggregates. We identified over 3770 protein groups, among which 3290 were significantly more abundant in isolates from aged mice compared with the samples derived from young mice (Fig. 3B, Table S1). We assume that low-level detection of peptides from the young spleens may represent reminiscence from intracellular deposits that we observed via TEM and which may be released from RPMs during splenocyte processing. We performed a functional enrichment analysis of the top 387 hits (log2 fold change >5) with DAVID and ShinyGO (Fig. 3B and C). We first noticed that although the identified proteins are enriched with components of several organelles, including lysosomes, ER, mitochondria, but also nucleus, and extracellular space, one of the top characteristics that they share is a disordered region in their amino acid sequence, a factor that may increase the risk of protein aggregation (Uversky, 2009). Consistently, the top hits are enriched in proteins related to neurodegeneration pathways, amyotrophic lateral sclerosis, Huntington’s disease, and heat shock protein binding, linking their origin to proteostasis defects (Medinas et al., 2017). For example, these include huntingtin (HTT) and ataxin 3 (ATXN3), the latter underlying spinocerebellar ataxia-3, both proteins prone to polyglutamine track expansion, and calpain (CAPN2), a protease implicated in neuronal cell death in neurodegeneration (Camins et al., 2006; Lieberman et al., 2019) (Fig. 3B). In agreement with the substantial iron load of the aggregates and the fact that iron promotes protein aggregation (Klang et al., 2014), we identified “metal-binding” as another significant functional enrichment. Furthermore, we observed that the top components of the aggregates include several proteins associated with immunoglobulin-like domains (Fig. 3B), such as 15 antibody chain fragments as well as 2 Fc receptors, proteins associated with complement activation and phagocytosis, thus linking the aggregates to the removal of defective cells, likely senescent RBCs or damaged RPMs. Finally, consistent with our data from EM imaging that cell death likely contributes to the formation of the aggregates (Fig. 3Aiv), their top components are enriched with apoptosis-related protein (e.g. BID, DFFA, DAXX, and MRCH1; Fig. 3B) and those linked to the response to stress. To explore how limited dietary iron content during aging affects the formation of the splenic iron-rich deposits, we conducted TMT-based proteomic quantification of their composition. Out of 942 detected protein groups, 50 hits were significantly more abundant in magnetically-isolated aggregates from the aged mice than mice fed an iron-reduced diet (Fig. 3D, Table S2). Functional enrichment analysis revealed that proteins related to phagosomes, response to stress, or necroptosis distinguish the rescued mice from those aged on a standard diet (Fig. 3E). Interestingly, however, this analysis identified components of lysosomes as by far the most overrepresented hits (Fig. 3D and E), suggesting that alleviation of the lysosomal defects (Fig. 1K) likely contributes to less pronounced aggregates formation in iron-reduced mice. Furthermore, 50 hits include both H and L ferritin as being significantly increased in aggregates from aged mice on a standard diet (Fig. 3D), and we now detected “ferric iron-binding” as an overrepresented characteristic within this group (Fig. 3E). Taken together, our data suggest that the splenic heme- and iron-rich deposits are composed of a wide spectrum of aggregation-prone protein debris that likely emerges from damaged RPMs.

**Figure 3.**
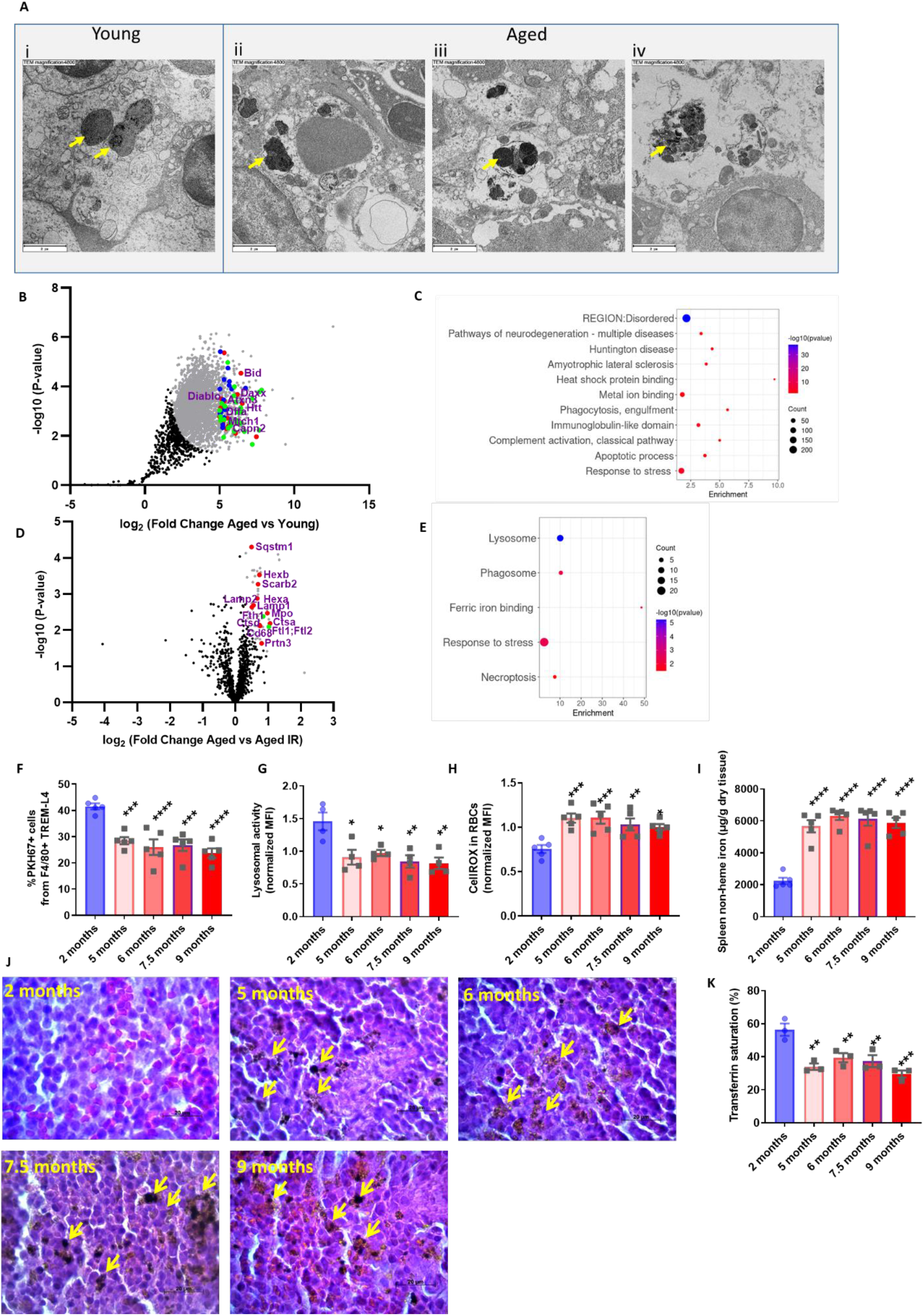
Splenic age–triggered iron deposits are rich in aggregation-prone protein debris that likely emerges from dysfunctional and damaged RPMs. (A) Ultrastructural analyses of the spleen red pulp sections by transmission electron microscopy. Arrows indicate dark-colored dense deposits. (B) Volcano plot illustrating 3290 protein groups (in grey) that are significantly more abundant in magnetically-sorted, cell-free aggregates derived from aged *versus* young spleen. Based on the functional enrichment analyses, the red color denotes proteins linked with “pathways of neurodegeneration”, green - those associated with an “apoptotic process” and blue - those related to the “immunoglobulin-like domain” category. (C) Enriched functional categories among top 387 protein hits that are more abundant in magnetically-sorted, cell-free aggregates derived from aged *versus* young spleen. (D) Volcano plot illustrates protein groups (in grey) that are significantly more abundant in magnetically-sorted, cell-free aggregates derived from aged *versus* aged IR spleen. Based on the functional enrichment analyses, the red color denotes proteins linked with “lysosome” and green – H and L ferritin. (E) Enriched functional categories among 50 protein hits that are more abundant in magnetically-sorted, cell-free aggregates derived from aged *versus* aged IR spleen. (F) RBC clearance capacity of RPMs in mice at the indicated age was determined by measuring the percentage of RPMs that phagocytosed transfused PKH67-labeled temperature-stressed RBCs. (G) The lysosomal activity of RPMs derived from mice at the indicated age was determined using a dedicated fluorescent probe and flow cytometry. (H) The cytosolic ROS levels in RPMs derived from mice at the indicated age were assessed by determining CellROX Deep Red fluorescence intensity with flow cytometry. (I) Spleen non-heme iron content was determined in mice at the indicated age. (J) Hematoxylin/eosin staining of the splenic red pulp in young, aged, and aged IR mice. Arrows indicate extracellular brown protein aggregates. (K) Plasma transferrin saturation was determined in mice at the indicated age. Each dot represents one mouse. Data are represented as mean ± SEM. Statistical significance among the three or more groups was determined by One-Way ANOVA test with Tukey’s or Dunnett’s Multiple Comparison test. *p<=0.05, **p<=0.01, ***p<=0.001 and ****p<=0.0001

To further explore links between splenic iron deposition and RPMs’ functions, we monitored in a time-wise manner how the formation of iron-rich aggregates corresponds to the iron-recycling performance of RPMs. We found that RPMs reduce their EP and lysosomal degradation capacity early during age progression (Fig. 3F and G). Correspondingly to the EP drop in mice at 5 months of age, we observed that at the same time point splenic RBCs exhibit the senescence marker, increased ROS levels (Fig. 3H). Interestingly, these changes reflect the kinetics of splenic non-heme iron loading (Fig. 3I). Likewise, we observed that the extracellular splenic protein deposits appear in 5-months-old mice, and their abundance increases during aging (Fig. 3J). Since these deposits are insoluble, we propose that their formation limits the bioavailability of iron for further systemic utilization. In support of this possibility, we noticed that the drop in transferrin saturation during aging coincides with RPMs’ failure and the appearance of splenic aggregates (Fig. 3K). In sum, our data support the model that defects in the iron-recycling capacity of RPMs, along with their damage, drive iron retention in the spleen in the form of non-bioavailable protein-rich deposits.

### Iron-recycling dysfunction during aging involves RPM ferroptosis

Increased iron burden in RPMs was shown previously to drive ferroptotic cell death upon acute transfusion of damaged RBCs (Youssef et al., 2018). Since the TEM imaging revealed damaged iron-loaded RPMs in aged spleens, we speculated that RPMs may undergo spontaneous ferroptosis during aging. In agreement with this hypothesis, both labile iron, a factor that promotes ferroptosis, and lipid peroxidation, a ferroptosis marker (Dixon et al., 2012; Friedmann Angeli et al., 2014) increase in RPMs during aging following a similar pattern as their functional impairment and the formation of iron-rich splenic protein aggregates (Fig. 4A and B). Importantly, we observed that feeding mice an IR diet during aging diminishes lipid peroxidation to the levels characteristic of young mice (Fig. 4C), similarly to what we observed for the LIP build-up (Fig. 1G). In agreement with the TEM data, we also detected a lower representation of RPMs in spleens of aged mice, a phenotype that likewise was alleviated by limited dietary iron content (Fig. 4D). Earlier studies reported that ferroptotic cells are characterized by reduced mitochondria size but not mitochondrial oxidative stress (Chen et al., 2021; Dixon et al., 2012; Friedmann Angeli et al., 2014). Others demonstrated that cells that undergo ferroptosis exhibit diminished mitochondrial membrane potential and a drop in cellular ATP levels (Neitemeier et al., 2017). Consistent with these data, we failed to detect augmented ROS levels in mitochondria of aged RPMs (Fig. 4E). Instead, we observed that mitochondria mass (measured with the MitoTracker probe, Fig. 4F) as well as mitochondrial activity (determined by the TMRE probe that quantifies mitochondrial membrane potential, Fig. 4G) decrease in RPMs of aged mice, and both these phenotypes are alleviated by an IR diet. This corresponds to higher ATP levels in FACS-sorted RPMs derived from aged mice fed an IR diet compared to the standard diet (Fig. 4H). Previous ultrastructural studies showed that mitochondria of ferroptotic cells are hallmarked with a reduced size and fewer cristae, increased membrane density, and some appear swollen (Chen et al., 2021; Dixon et al., 2012; Friedmann Angeli et al., 2014). Consistently, our TEM imaging showed that in contrast to the RPMs of young mice, those from aged animals exhibit the above mitochondrial defects and some seem disintegrated (Fig. 4I). Mitochondria from mice fed an IR diet were small and dense, but they display a lesser degree of swelling and damage. Taken together, our data suggest that aging is hallmarked by both impaired iron-recycling functions of RPMs and their depletion likely via ferroptosis, both defects that we found alleviated by the reduction of dietary iron content.

**Figure 4.**
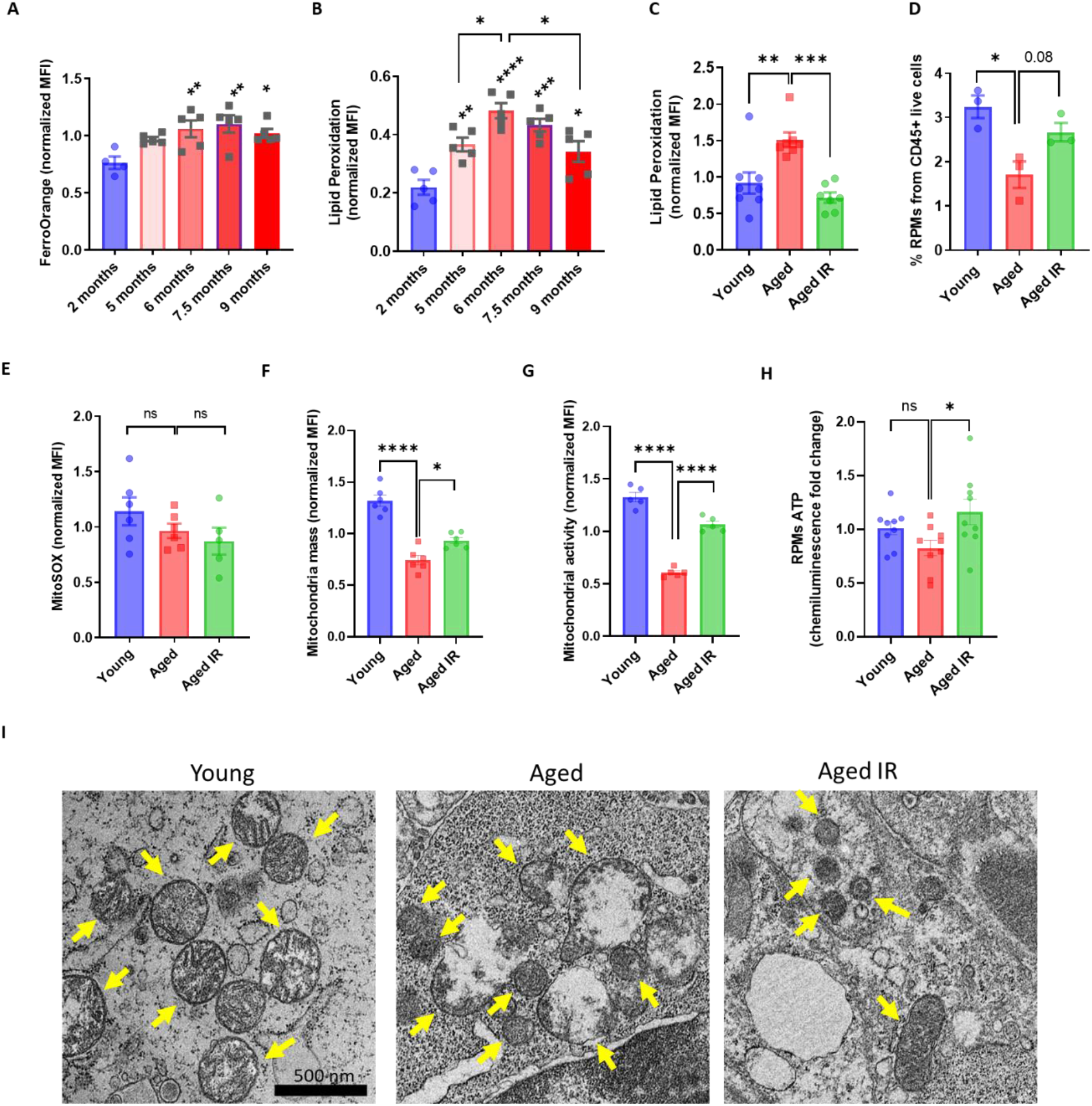
Iron-recycling dysfunction during aging involves RPM ferroptosis. (A) Cytosolic ferrous iron (Fe2+) content in RPMs isolated from mice at the indicated age was measured using FerroOrange and flow cytometry. (B) Lipid peroxidation was determined in RPMs derived from mice at the indicated age using a dedicated ratiometric fluorescent probe and flow cytometry. (C) Lipid peroxidation was determined in RPMs derived from young, aged, and aged IR mice using a dedicated ratiometric fluorescent probe and flow cytometry. (D) Shown is the percentage of RPMs from CD45+ live cells present in the spleen of young, aged, and aged IR mice. (E) Mitochondrial ROS levels in RPMs derived from young, aged, and aged IR mice were determined using the MitoSOX probe and flow cytometry. (F) The mitochondrial mass and (G) mitochondrial membrane potential in RPMs isolated from young, aged, and aged IR mice were measured by staining with MitoTracker Green and TMRE, respectively, and analyzed by flow cytometry. (H) Levels of ATP in FACS-sorted RPMs isolated from young, aged, and aged IR. (I) Ultrastructural analyses of mitochondrial morphology in spleen red pulp sections obtained from young, aged, and aged IR mice. Yellow arrows indicate mitochondria. Each dot represents one mouse. Data are represented as mean ± SEM. Statistical significance among the three or more groups was determined by One-Way ANOVA test with Tukey’s or Dunnett’s Multiple Comparison test. *p<=0.05, **p<=0.01, ***p<=0.001 and ****p<=0.0001

### Iron loading, but not oxidative stress or diminished mitochondrial function of RPMs suppresses the iron-recycling capacity

The contribution of iron deposition to age-related deterioration of cellular functions is believed to be primarily mediated by the pro-oxidative properties of labile iron (Xu et al., 2008). We observed a clear indication of oxidative stress in aging RPMs (Fig. 1I) that was rescued by the IR diet. Next, we aimed to verify whether the increased ROS levels solely contribute to diminished phagocytic and cellular functions of RPMs during aging. To this end, we supplemented aging mice with antioxidant N-Acetyl-L-cysteine (NAC), which was previously shown to revert aging-related physiological changes (Berman et al., 2011; Ma et al., 2016). With this strategy, we successfully reduced oxidative stress in aged RPMs (Fig. 5A). However, we did not observe an improvement in the reduced rate of EP in RPMs isolated from NAC-supplemented mice compared to control aged mice (Fig. 5B). Likewise, the retention of senescent RBCs in the spleen (Fig. 5C), their local hemolysis (Fig. 5D), as well as the splenic iron loading (Fig. 5E), and the formation of the insoluble aggregates (Fig. 5F) were not rescued by the NAC administration. Lastly, the ferroptosis marker, lipid peroxidation was equally elevated in RPMs of aged mice regardless of the NAC treatment (Fig. 5G). These data suggest that the ability of RPMs to remove RBCs, and consequently iron and erythrocyte homeostasis in the spleen are likely affected directly by RPM iron loading rather than excessive ROS. To follow this possibility, we next employed a cellular model of bone marrow-derived macrophages, exposed to hemin and IL-33, two factors that drive RPMs differentiation (Haldar et al., 2014; Lu et al., 2020) and induce RPM-like transcriptional signatures (Lu et al., 2020). We noted that these cells that we termed “induced RPMs” (iRPMs) significantly decrease their EP activity upon iron overload with ferric ammonium citrate (FAC) in a manner that is partially rescued by the iron chelator DFO (Fig. 5H). Interestingly, adding the same dose of hemin as FAC did not significantly alter the rate of RBCs uptake by iRPMs, suggesting a low sensitivity of the phagocytic system of these cells to excessive heme. Furthermore, we observed that iron loading by FAC, but not hemin, suppresses lysosomal activity (Fig. 5I). We also found that mitochondria membrane potential was similarly diminished by FAC, hemin, and DFO alone, thus uncoupling the phagocytic and lysosomal activity of iRPMs from their mitochondria fitness (Fig. 5J). In sum, our data indicate a causative link between iron loading, a condition that reflects the physiological status of aging mouse RPMs, and the perturbation of the iron-recycling capacity of RPM-like cells.

**Figure 5.**
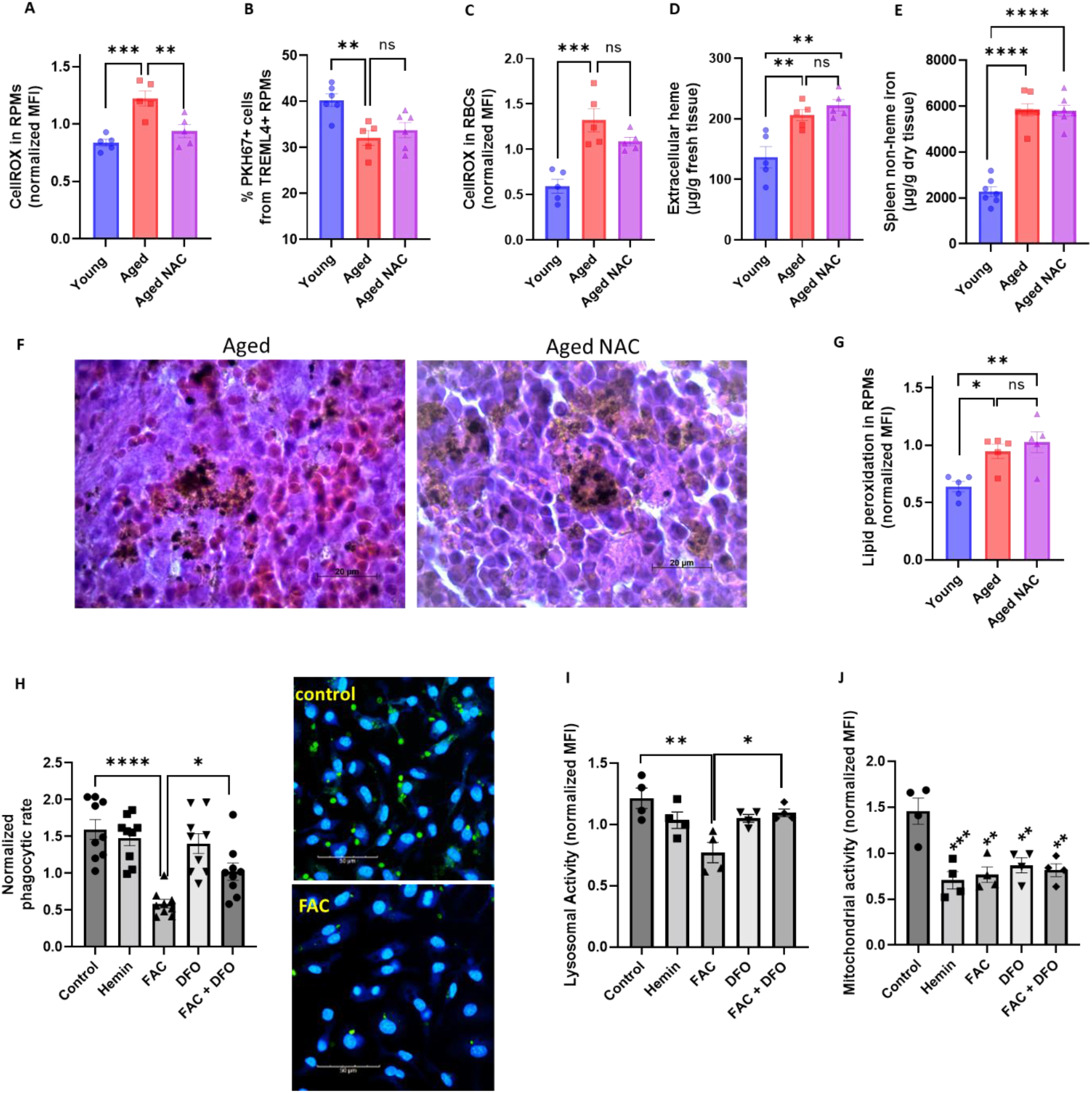
Iron loading, but not oxidative stress or diminished mitochondrial function of RPMs suppresses the iron-recycling capacity. (A) The cytosolic ROS levels in RPMs derived from young, aged, and aged NAC were assessed by determining CellROX Deep Red fluorescence intensity with flow cytometry. (B) RBC clearance capacity of RPMs derived from young, aged, and aged NAC mice was determined by measuring the percentage of RPMs that phagocytosed transfused PKH67-labeled temperature-stressed RBCs. (C) The cytosolic ROS levels in RBCs derived from the spleen of young, aged, and aged NAC mice were assessed by determining CellROX Deep Red fluorescence intensity with flow cytometry. (D) Extracellular heme levels were measured in the medium obtained after dissociation of spleens from young, aged, and aged NAC mice using Heme Assay Kit. (E) Spleen non-heme iron content was determined in young, aged, and aged NAC mice. (F) Hematoxylin and eosin staining of the splenic red pulp in young, aged, and aged IR mice. (G) Lipid peroxidation was determined in RPMs derived from young, aged, and aged NAC mice using a dedicated ratiometric fluorescent probe and flow cytometry. (H) Normalized phagocytic rate of PKH67-labeled temperature-stressed RBCs by cultured iRPMs. Cells were treated with FAC (50 μM, 24 h), hemin (50 μM, 24 h), or DFO (100 μM, 18 h) as indicated. Representative confocal microscopic images of erythrophagocytosis in FAC-treated iRPMs compared with control cells are shown on the right. (I) Lysosomal and (J) mitochondrial activity of cultured iRPMs were determined using dedicated fluorescent probes and flow cytometry. Cells were treated with FAC (50 μM, 24 h), hemin (50 μM, 24 h), or DFO (100 μM, 18 h) as indicated. Each dot represents one mouse or independent cell-based experiment. Data are represented as mean ± SEM. Statistical significance among the three or more groups was determined by One-Way ANOVA test with Tukey’s Multiple Comparison test. *p<=0.05, **p<=0.01, ***p<=0.001 and ****p<=0.0001

### Iron loading undermines the phagocytic activity of aged RPMs in concert with impaired heme metabolism and ER stress

Our data so far imply that iron accumulation in RPMs suppresses their iron-recycling functions. However, restoration of iron status in RPMs of aging mice by IR diet does not fully recover EP activity (Fig. 1J). To further understand the molecular mechanisms that influence phagocytic RPMs dysfunction during aging, we performed RNA sequencing of FACS-sorted RPMs from young and aged mice. We identified 570 differentially expressed genes, including 392 up- and 178 down-regulated transcripts (Fig. S7A). Interestingly, only 54 genes were found differentially regulated between mice fed a standard and IR diet (Fig. S7B), implying that functional changes between these groups are largely based on protein levels, activity and/or post-transcriptional changes. Functional enrichment analysis revealed that aged RPMs, regardless of the dietary regimen, are hallmarked by ER stress, unfolded protein response, and ER-associated degradation (ERAD) (Fig. 6A and S7C). This observation corroborates the finding that extracellular iron-rich protein aggregates, which appear, albeit to a lesser extent, in IR mice, are enriched in proteins that imply proteotoxic stress in aged RPMs (Fig. 3C). ERAD is a pathway for targeting misfolded ER proteins for cytoplasmic proteasomal degradation (Qi et al., 2017). Correspondingly, we identified a significant increase in proteasomal activity in aged RPMs, measured with a dedicated fluorescent probe that undergoes proteasomal cleavage (Fig. 6B). The IR diet rescued this phenomenon, suggesting links with iron/oxidative stress-related factors and potentially a compensatory response to the lysosomal-mediated proteostatic control (Fig. 1K). Furthermore, our RNA-seq data revealed that one of the transcripts that significantly decrease in aged RPMs independent of diet is *Hmox-1*, which encodes HO-1. We validated this result using intracellular staining and flow cytometry, and uncovered a marked decrease in HO-1 protein level in aged RPMs (Fig. 6C). Interestingly, this HO-1 drop seems to progress between 5 to 9 months of age (Fig. 6D), thus reflecting both diminished EP rate and lysosomal activity (Fig. 3F and G). Correspondingly, the blockage of HO-1 in iRPMs with zinc protoporphyrin (ZnPP) suppressed the EP rate (Fig. 6E) and, to a lesser extent, the capacity for lysosomal degradation (Fig. 6F). RPMs derived from aging mice show similar HO-1 levels and the degree of ER stress irrespective of diet and differ primarily in iron status (Fig. 1G and H). Therefore, we next tested how the combination of a relatively low dose of ZnPP (0.5 μM) and the ER stress inducer tunicamycin with a mild FAC exposure (10 μM) affects EP *in cellulo*. Whereas ER stress alone did not significantly affect EP, we uncovered that FAC treatment elicited an additive effect with combined exposure to ZnPP and tunicamycin, but not with ZnPP alone (Fig. 6G). Finally, we explored why ZnPP, but not hemin alone, suppresses EP. We speculated that possibly another co-product of HO-1 enzymatic activity, such as CO or biliverdin (Gozzelino et al., 2010), counterbalances the suppressive effect of iron release after EP. Thus, we tested whether the effect of ZnPP on EP can be rescued by supplementation of cells with CO donor CORMA1 or biliverdin. Interestingly, we found that CORMA1, but not biliverdin, rescues EP rate under ZnPP exposure in iRPMs (Fig. 6H and I), implying that CO acts as a modulator of EP activity in RPMs. Collectively, our data imply that iron retention in RPMs suppresses EP in synergy with ER stress, HO-1 loss, and possibly CO deficiency, but likely represent the key factor that determines their RBC clearance potential.

**Figure 6.**
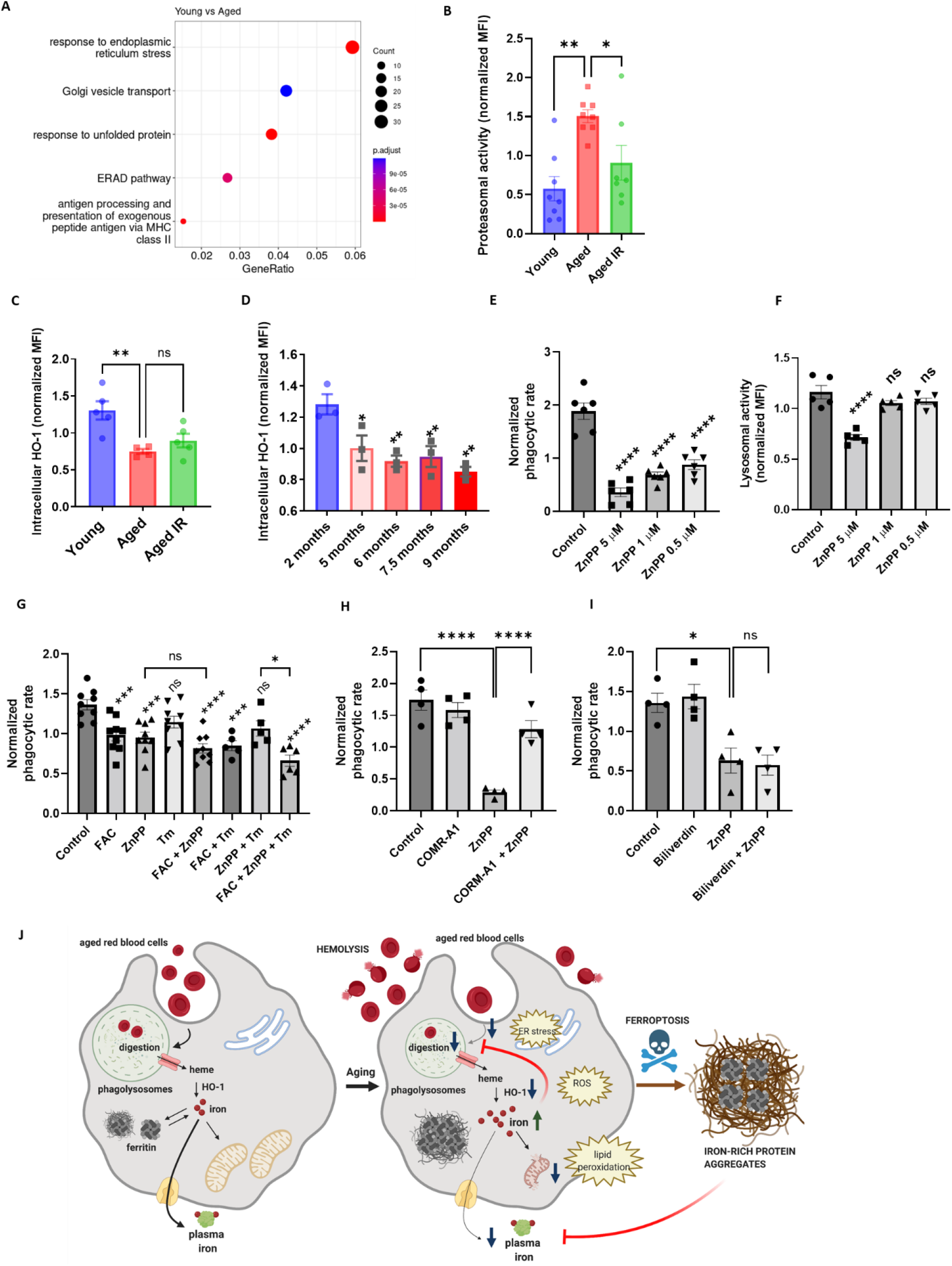
Iron loading undermines the phagocytic activity of aged RPMs in concert with impaired heme metabolism and ER stress. (A) Enriched functional categories among differentially regulated genes in FACS-sorted RPMs derived from young *versus* aged mice (identified by RNA-seq). (B) Proteasomal activity measured in RPMs isolated from young, aged, and aged IR mice using a dedicated fluorescent probe and flow cytometry. (C) Intracellular HO-1 protein levels in RPMs isolated from young, aged, and aged IR mice. (D) Intracellular HO-1 protein levels in RPMs isolated from mice at the indicated age. (E) Normalized phagocytic rate of PKH67-labeled temperature-stressed RBCs and (F) normalized lysosomal activity by cultured iRPMs. Cells were treated with indicated concentrations of the HO-1 inhibitor ZnPP for 24 h. (G) Normalized phagocytic rate of PKH67-labeled temperature-stressed RBCs by cultured iRPMs. Cells were treated with indicated low concentrations of ZnPP (0.5 μM) and FAC (10 μM) and with ER stress inducer Tunicamycin (Tm; 2.5 μM) for 24 h. (H and I) Normalized phagocytic rate of PKH67-labeled temperature-stressed RBCs by cultured iRPMs. Cells were treated with ZnPP (5 μM) and (H) CORMA1 (50 μM) or (I) Biliverdin (50 μM). (J) The proposed model of aging-triggered impairment of the iron-recycling functions of RPMs. Each dot represents one mouse or independent cell-based experiment. Data are represented as mean ± SEM. Statistical significance among the three or more groups was determined by One-Way ANOVA test with Dunnett’s or Tukey’s Multiple Comparison test. *p<=0.05, **p<=0.01, ***p<=0.001 and ****p<=0.0001

## Discussion

RPMs play an essential role in preventing the release of hemoglobin from RBCs and maintaining the internal recycling of the body iron pool (Slusarczyk and Mleczko-Sanecka, 2021). To date, impairments of their functions have been reported in genetically modified mice, including models with an almost complete or partial loss of RPMs due to the absence of genes that control RPMs differentiation, such as *Pparg, Spi-c, Il-33*, or its receptor *Il1rl1* (Kohyama et al., 2009; Lu et al., 2020; Okreglicka et al., 2021). Here, for the first time, we provide evidence that the defective RPMs’ capacity for RBC clearance and degradation together with ferroptotic RPMs depletion accompanies physiological aging, promoting local RBC dyshomeostasis and the formation of protein-rich extracellular iron aggregates in the spleen (Fig. 6J). We demonstrate that these phenomena can be mitigated by reducing dietary iron availability.

Since the unique architecture of the spleen confers a filter for verifying the biomechanical integrity of RBCs (Mebius and Kraal, 2005; Slusarczyk and Mleczko-Sanecka, 2021), we observed only local but not systemic dyshomeostasis of RBCs during aging. Most importantly, we found that splenic RBCs of aged mice show ROS accumulation, a characteristic feature of senescent RBCs, and a risk factor for enhanced hemolysis (Zhang et al., 2018). Recent insights suggest that iron recycling in the spleen is driven by an equilibrium between phagocytosis and natural hemolysis of RBCs (Klei et al., 2020). Our study implies that this balance is likely shifted towards the extracellular destruction of senescent RBCs in aging. It remains to be investigated whether the exposure of the splenic microenvironment to excessive heme due to the enhanced breakdown of RBCs may affect the function of other cells that traffic and reside in the spleen, such as lymphocytes.

The present study provides, what we believe, is new evidence for the origin and nature of iron deposits in the aging spleen that also hallmark genetic mouse model of impaired iron-recycling (Kohyama et al., 2009; Lu et al., 2020; Okreglicka et al., 2021). We found that they are rich in iron and heme, suggesting that they may still be derived, likely not directly, from erythrocytes and contain many proteins (>3000) of different subcellular origins. Since the aggregates are enriched in proteins related to phagocytic removal of antibody and complement opsonized cargos and in lysosomal components, they may be a reminiscence of phagocytic and lytic events of RPMs. However, our observations imply that two driving forces underlie the formation of these deposits: proteostasis defects and RPMs damage.

The longevity of RPMs (Hashimoto et al., 2013; Liu et al., 2019) puts pressure on the cell to handle the constant flux of iron, a factor that increases protein insolubility (Klang et al., 2014), and RBC-derived proteins. These may increase the risk for the build-up of proteotoxic elements in RPMs. Consistently, we found a clear transcriptional signature for ER stress and unfolded protein response in aged RPMs. Interestingly, it was also reported that phagocytosis triggers ER stress in macrophages (Kim et al., 2018), providing another explanation of how ER homeostasis may be affected in constitutive phagocytes such as RPMs. The aggregates that emerge in the aging spleen are chiefly composed of aggregation-prone proteins that contain disordered regions and are associated with different pathways of neurodegeneration. It has been reported previously that excessive iron in the brain promotes the progression of neurodegenerative disorders, such as Alzheimer’s disease (Liu et al., 2018; Raven et al., 2013). Interestingly, iron deposits were shown to spacially colocalize with β-amyloid plaques (Ndayisaba et al., 2019; van Bergen et al., 2016). Our findings provide a new link between splenic and brain protein aggregation that is known to underlie cognitive impairment. Future studies would be required to investigate how these protein deposits may affect other (e.g. immune) functions in the spleen. Moreover, our work suggests that in addition to various iron chelation approaches (Liu et al., 2018), long-term reduction of dietary iron intake may limit protein aggregation in the brain and thus represent an alternative for dementia prevention.

Second, our data imply that RPMs undergo spontaneous ferroptotic cell death, a response that thus far was mainly described in genetic models, e.g. with disturbed detoxification of lipid peroxides due to inducible or cell type-specific loss of *Gpx4* (Friedmann Angeli et al., 2014; Matsushita et al., 2015) or in various pathologies, such as ischemia-reperfusion injury, myocardial infarction or neurodegeneration (Park et al., 2019; Yan et al., 2021). In the latter context, iron-rich activated microglial cells were shown to localize to β-amyloid deposits and undergo cell death upon plaques phagocytosis (Baik et al., 2016; Kenkhuis et al., 2021). This may partially resemble the phenomenon that we observe in the aging spleen, opening a possibility that functional RPMs may attempt to phagocytose the extracellular protein deposits, which may lead to their damage. Notably, the excessive clearance of damaged RBCs in mouse models of blood transfusion, hemolytic anemia, or the anemia of inflammation induced by heat-killed *Brucella abortus* was previously shown to cause RPMs loss, likely via ferroptosis (Haldar et al., 2014; Theurl et al., 2016; Youssef et al., 2018). Whereas the splenic niche may be replenished to some extent via monocyte recruitment and/or resident RPMs proliferation (Haldar et al., 2014; Youssef et al., 2018) our data imply that RPM depletion may lead to the irreversible formation of insoluble iron-rich splenic protein aggregates, a possibility that would require further investigations. Premature clearance of defective RBCs also hallmarks β-thalassemia (Slusarczyk and Mleczko-Sanecka, 2021). To the best of our knowledge, RPM loss was not extensively studied in a mouse model of thalassemia. However, interestingly, thalassemic mice display enhanced splenic Perls’-stained deposits that appear extracellular and are not amenable to iron chelation therapy (Sanyear et al., 2020; Vadolas et al., 2021), possibly resembling those identified by the present study.

Of note, our findings that the aggregates are enriched in apoptosis-related proteins are not in conflict with the proposed involvement of ferroptotic cell death. One of such proteins, for example, is BID, an apoptosis agonist that mediates mitochondrial defects in ferroptotic neurons, and also was shown to be required for the execution of ferroptosis itself (Neitemeier et al., 2017). Moreover, several studies showed that the induction of ferroptosis in mice, for example in the context of acute renal injury, is hallmarked by DNA fragmentation detected by the TUNEL staining (Friedmann Angeli et al., 2014), a typical apoptosis marker.

Genetic disruption of *Hrg-1*, a transporter that delivers RBC-derived heme to the cytoplasm, leads to the formation of heme aggregate hemozoin inside enlarged phagolysosomes of RPMs (Pek et al., 2019). The large size, extracellular localization, and high content of proteins distinguish age-triggered splenic aggregates from hemozoin. However, we cannot exclude the possibility that this polymer-like structure may expand during aging and contribute to the formation of splenic protein-rich iron deposits.

Erythrocyte removal by iron recycling macrophages is considered an essentially constitutive process. Nevertheless, recent advances revealed that it remains under the control of calcium signaling and can be modulated by inflammatory signals (Bennett et al., 2019; Ma et al., 2021). Interestingly, it was unknown if the RBCs uptake rates are regulated by iron availability *per se*. Our study identified three factors that affect the intensity of EP, the iron content, the activity of HO-1, and, to less extent, ER stress. Among them, labile and total iron accumulation in RPMs, independently of ROS generation, emerged as a major driver of EP suppression during aging. We also demonstrated that iron loading decreases the lysosomal activity of iRPMs and that lysosomal activity in RPMs *in vivo* closely reflects their iron status. Consistently, excessive iron was shown by others to promote ROS-triggered lysosomal membrane permeabilization (Kurz et al., 2011) and to inhibit phagosome fusion with lysosomes (Kelley and Schorey, 2003). Our data assign physiological context to this previous knowledge, as RPMs constitutively rely on lysosomal activity to exert their functions. The underlying mechanisms for excessive iron deposition in aged RPMs likely include reduced FPN protein levels and deficient functional H ferritin that detoxifies redox-active ferrous iron (Mleczko-Sanecka and Silvestri, 2021). It is also plausible that, like neurons, aged RPMs may also show low capacity for heme synthesis (Atamna et al., 2002), which may lead to defective iron utilization in the mitochondria and increased cytoplasmic iron burden. Mitochondrial damage that we identified in aged RPMs may support this possibility.

Another medically relevant condition hallmarked by iron retention in RPMs is inflammation (Guida et al., 2015; Kim et al., 2014). It remains to be investigated whether under such settings iron accumulation contributes to the iron-recycling capacity and possibly RPMs loss. Given that the net outcome in young mice is EP stimulation (Bennett et al., 2019; Bian et al., 2016; Delaby et al., 2012), it is plausible that inflammatory signals override the suppressive effect of iron. Our study also suggests that aged mice characterized by defective RPMs may not potentiate EP rate in response to inflammation. Since this phenomenon was proposed to activate stress erythropoiesis (Bennett et al., 2019), age progression may impair this adaptive response and impede recovery from inflammation-triggered anemia.

The heme catabolizing enzyme HO-1 is well characterized for its cytoprotective, anti-oxidative, and anti-inflammatory function, mainly attributed to its products, CO and biliverdin (Gozzelino et al., 2010). More recently, HO-1 activity was proposed to prevent cellular senescence in endothelial cells, fibroblasts or macrophages, via decreasing ROS levels, improving mitochondria function, or protecting from DNA damage (Even et al., 2018; Hedblom et al., 2019; Luo et al., 2018; Suliman et al., 2017). Iron-recycling RPMs are key cells where HO-1 exerts its enzymatic function (Vijayan et al., 2018) and complete HO-1 deletion in mice leads to RPMs death due to heme-driven toxicity (Kovtunovych et al., 2010). Our observation that HO-1 levels decrease in these cells during aging is thus of high physiological significance. As aged RPMs show some hallmarks of senescence linked to HO-1 deficiency, such as defective mitochondria or excessive ROS, they do not exhibit proinflammatory gene expression signatures. Instead, our RNA-seq data reveal rather an anti-inflammatory transcriptional pattern, exemplified by the downregulation of genes encoding for MHC class II complexes and the induction of *Il-10*. Future research may address if such skewing of aged RPMs may contribute to immunosenescence that accompanies aging (Nikolich-Zugich, 2018). Interestingly, the expansion of iron-recycling KCs that we observed in aged mice may exert further immunosuppressive functions, as shown previously for the liver macrophages exposed to erythrolytic stress (Olonisakin et al., 2020; Pfefferle et al., 2020).

The observation that HO-1 inhibition suppresses EP in RPM-like cells may be underlain by the inhibitory effect of heme overload on general phagocytosis in macrophages (Martins et al., 2016). However, unlike standard BMDMs (data not shown), heme-supplemented iRPMs failed to modulate EP in response to exposure to excessive hemin, implying efficient heme catabolism via HO-1 in this model. We further provide evidence that another product of HO-1, CO, restores the EP capacity of cells subjected to HO-1 suppression, thus emerging as an inducer of RBCs uptake. These findings are in agreement with the previous work, reporting that CO administration protects HO-1 knock-out mice from sepsis-induced lethality via stimulation of bacterial phagocytosis (Chung et al., 2008). Whether deficient CO levels in RPMs *in vivo* contribute to EP suppression requires further investigation. Nevertheless, in support of this possibility, our RNA-seq data show that enzymes involved in the pentose phosphate pathway, such *Tkt, Taldo* or *G6pd*, transcriptional targets of CO-mediated regulation (Bories et al., 2020), seem downregulated in aged RPMs.

Collectively, our work demonstrates that age-triggered and iron-dependent functional defects of RPMs, specialized cells that supply iron to the organism, emerge during early aging and disrupt internal iron turnover (Fig. 6J). We propose that the build-up of un-degradable iron and protein-rich particles, in concert with increased hepcidin levels, limits plasma iron availability during aging. Future studies may be needed to design strategies for complete “rejuvenation” of the RPMs’ phagocytic activity or proteostatic homeostasis, to completely alleviate the excessive splenic hemolysis, the formation of iron-rich aggregates, and hence possibly fully restore plasma iron availability.

## Materials and Methods

### Mice

Female C57BL/6J mice were used for all the experiments and were maintained in specific pathogen-free conditions at the Experimental Medicine Centre (Bialystok, Poland). Starting with the age of 4 weeks mice were fed a diet with a standard iron content (200 mg/kg, SAFE #U8958v0177; for Young and Aged), as described before (Kautz et al., 2008; Pagani et al., 2011) or reduced iron content [25 mg/kg; SAFE #U8958v0294; for Aged iron reduced (IR)]. Diets were from SAFE (Augy, France). Mice were analyzed at 8-10 weeks (Young) and 10-11 months (Aged and Aged IR) of age. For supplementation with N-Acetyl-L-cysteine (NAC), aging mice (Aged NAC) were supplied with NAC dissolved in drinking water (2 g/L) from 8 weeks of age until 10-11 months of age. Mice were delivered to the local facility at the IIMCB and sacrificed after short acclimatization, or subjected to additional procedures, if applicable. All procedures were approved by the local ethical communities in Olsztyn and Warsaw (II LKE) (decisions: WAW2/015/2019; WAW2/149/2019; WAW2/026/2020; WAW2/149/2020).

### Preparation of single-cell suspension from mouse organs

**Bone marrow** cells were harvested by flushing the femur and tibia using a 25G needle and sterile HBSS medium (Gibco, 14025092). Cells were centrifuged at 600g for 10 min at 4°C. **The spleen** was excised and mashed through a 70μm strainer (pluriSelect, 43-50070-51). For FACS and magnetic sorting, the spleen was additionally digested in HBSS medium containing 1 mg/ml Collagenase D (Roche, 11088882001) and 50 U/ml DNase I for 30 min at 37°C. After that cells were washed with cold HBSS and centrifuged at 600g for 10 min at 4°C. **The liver** was extracted and perfused using Liver Perfusion Medium (Gibco, 17701038). Next, the organ was minced and digested in Liver Digest Medium (Gibco, 17703034) for 30 min at 37°C. After that liver was pressed through a 70μm strainer in the presence of HBSS. Cells were centrifuged at 50g for 3 min and the pellet was discarded (hepatocytes). The supernatant was centrifuged at 700g for 15 min at 4°C. Pellet was resuspended in 5mL of PBS containing 0,5% BSA and 5mL of 50% Percoll (Cytiva, 17-0891-01) diluted in PBS. The suspension was centrifuged at 700g for 30 min at 20°C. **Blood** was collected to a heparin-coated tube *via* heart puncture. Cells were washed with HBSS medium and centrifuged at 400g for 10 min at 4°C. For **peritoneal cells** isolation, peritoneal cavities were washed with HBSS medium, and the peritoneal fluid was aspirated and filtered through 70μm strainer, and cells were centrifuged at 600g for 5 min at 4°C.

RBCs present in a single-cell suspension were lysed using 1X RBC Lysis buffer (BioLegend, 420302) for 3 min at 4°C. This step was omitted for analyses of erythroid cells and RBCs. Next, cells were washed with HBSS and centrifuged at 600g for 10 min at 4°C. Pellet was prepared for further functional assays and/or labeling with antibodies.

### Generation of iRPMs and treatments

RPM-like cells (iRPMs) represent modified cultures of bone-marrow-derived macrophages. Mononuclear cells were obtained from femurs and tibias of female 2-4-months-old C57BL/6J mice. Sterile harvested cells were cultured at 37°C in 5% CO_2_ at 0.5 × 10^6^/1mL concentration in RPMI-1640 (Sigma-Aldrich, R2405) supplemented with 10% FBS (Cytiva, SV30160.03), 1X Penicillin-Streptomycin (Gibco, 15140122), and 20 ng/mL macrophage colony-stimulating factor (BioLegend, 576406). On the 4^th^ and 6^th^ day medium was changed to fresh, supplemented with 20 μM of hemin (Sigma-Aldrich, 51280) and 10 ng/mL of IL-33 (BioLegend, 580506). Assays were performed on the 8^th^ day.

For treatments, ferric ammonium citrate (FAC, Sigma-Aldrich, F5879), CORM-A1 (Sigma-Aldrich, SML0315), and deferoxamine (Sigma-Aldrich, D9533) were diluted in sterile ddH_2_O. Zinc (II) Protoporphyrin IX (Sigma-Aldrich, 691550) and tunicamycin (Sigma-Aldrich, T7765) were diluted in anhydrous DMSO. Hemin (Sigma-Aldrich, 51280) solution was prepared with 0.15M NaCl containing 10% NH4OH. All reagents after dilution were filtered through a 0,22μm filter and stored in −20°C except ferric ammonium citrate, which was always freshly prepared. Concentrations and time of treatments are indicated in the descriptions of the figures.

### Flow cytometric analysis and cell sorting

Cell suspensions of spleens and livers and iRPMs (~ 1 × 10^7^) were stained with LIVE/DEAD Fixable Aqua/Violet (Invitrogen, L34966/L34964) as per the manufacturer’s instructions to identify dead cells. After extensive washing, the cells were incubated with Fc block in a dilution of 1:100 in FACS buffer for 10 min at 4°C. Cells were then stained with fluorophore-conjugated antibodies, dilution of 1:100 to 1:400, depending on the titration, in FACS buffer for 30 min at 4°C. Cells were washed thoroughly with FACS buffer and subjected to flow cytometry analysis. For analysis of splenic RPM populations, the following surface antibodies were used: CD45 (Biolegend, 30-F11), CD45R/B220 (Biolegend, RA3-6B2), F4/80 (Biolegend, BM8), CD11b (Biolegend, M1/70) and TREML4 (Biolegend, 16E5). TER-119 (Biolegend) and CD71 (Biolegend, RI7217) were included for erythroid cell analysis. For analysis of liver Kupffer cell and iRPM populations, the following surface antibodies were used: CD45 (Biolegend, 30-F11), F4/80 (Biolegend, BM8) and CD11b (Biolegend, M1/70). Detection of FPN was performed with a non-commercial antibody that recognizes the extracellular loop of mouse FPN [rat monoclonal, Amgen, clone 1C7 (Sangkhae et al., 2019); directly conjugated with Alexa Fluor 488 Labeling Kit (Abcam, ab236553)]. For analysis of RBCs, the following surface antibodies were used: CD45 (Biolegend, 30-F11), TER-119 (Biolegend) and CD71 (Biolegend, RI7217). For analysis of peritoneal macrophages, the following surface antibodies were used: CD45 (Biolegend, 30-F11), F4/80 (Biolegend, BM8), MHCII (Biolegend, M5/114.15.2) and CD11b (Biolegend, M1/70). Events were either acquired on Aria II (BD Biosciences) or CytoFLEX (Beckman Coulter) and were analyzed with FlowJo or CytExpert, respectively. For RNA sequencing (RNA-seq) and ATP levels quantification, RPMs were sorted into Trizol-LS (Invitrogen, 10296010) or assay buffer, respectively, using an Aria II cell sorter (BD Biosciences) with an 85 μm nozzle.

### Functional Assays and detection of intracellular ferrous iron

**Intracellular ROS** (APC channel) levels were determined by using CellROX Deep Red Reagent (Invitrogen, C10422) fluorescence according to the manufacturer’s instructions. **Lipid peroxidation** (FITC vs PE channels) was determined with the Lipid Peroxidation Assay Kit (Abcam, ab243377) according to the manufacturer’s instructions. **Mitochondria-associated ROS** (PE channel) levels were measured with MitoSOX Red (Invitrogen, M36008) at 2.5 μM for 30 min at 37 °C. **Lysosomal activity** (FITC channel) was determined by using Lysosomal Intracellular Activity Assay Kit (Biovision, K448) according to the manufacturer’s instructions. **Proteasomal activity** (FITC channel) was determined using Me4BodipyFL-Ahx3Leu3VS fluorescent proteasome activity probe (R&D Systems, I-190) at 2 μM for 1 hour at 37°C. **Mitochondria activity** (membrane potential, PE channel) was measured using a tetramethylrhodamine ethyl ester (TMRE) fluorescent probe (Sigma-Aldrich, 87917) at 400 nM for 30 min at 37 °C. **Mitochondrial mass** (FITC/PE channel) was measured by fluorescence levels upon staining with MitoTracker Green (Invitrogen, M7514) at 100 nM for 30 min at 37 °C. The cells were subsequently stained with cell surface markers.

The content of **intracellular ferrous iron** (PE channel) (Fe^2+^) was measured using FerroOrange (DojinD) via flow cytometric analysis. Briefly, surface-stained cells were incubated with 1 μM FerroOrange in HBSS for 30 min at 37 °C, and analyzed directly via flow cytometry without further washing.

For **intracellular staining**, surface-stained cells were first fixed with 4% PFA and permeabilized with 0.5% Triton-X in PBS. The cells were then stained with respective primary antibodies for 1 h at 4°C followed by 30 min staining with Alexa Fluor 488- or Alexa Fluor 647-conjugated secondary anti-Rabbit IgG (1:1000 Thermo Fisher, A-21206). The following rabbit anti-mouse intracellular primary antibodies were used: Ferritin Heavy Chain (FTH1, Cell Signaling Technology, 3998), Ferritin Light Chain (FTL, Abcam, ab69090) and HO-1 polyclonal antibody (ENZO, ADI-SPA-896).

The geometric mean fluorescence intensities (MFI) corresponding to the probes/target protein levels were determined by flow cytometry acquired on Aria II (BD Biosciences) or CytoFLEX (Beckman Coulter) and were analyzed with FlowJo or CytExpert, respectively. For the probes that have emissions in PE, TREML4 was excluded, and RPMs were gated as F4/80-high CD11b-dim. For quantifications, MFI of the adequate fluorescence minus one (FMO) controls were subtracted from samples MFI, and data were further normalized.

### ATP levels

Cellular ATP levels in FACS-sorted RPMs (10 000 cells/sample) were determined using ATP Fluorometric Assay Kit (Sigma, MAK190), as per the manufacturer’s instructions.

### Magnetic sorting of RPMs and isolation of extracellular iron-containing aggregates

60 x 10^6^ of mouse splenocytes were incubated for 15 min at 4°C in PBS containing 5% Normal Rat Serum (Thermo Scientific, 10710C) and anti-CD16/32 (BioLegend, 101320) antibody in 5mL round bottom tube. Afterward, cells were labeled with anti-F4/80 (APC, BioLegend, 123116), anti-Ly-6G/Ly-6C (Gr-1) (Biotin, BioLegend, 108403), anti-CD3 (Biotin, BioLegend, 100243), anti-mouse Ly-6C (Biotin, BioLegend, 128003) and anti-B220 (Biotin, BioLegend, 103204) for 20 min in 4°C in dark. Next, cells were washed with cold PBS containing 2 mM EDTA, 0.5% BSA (hereafter referred to as “sorting buffer”), and centrifuged at 600g for 10 min. Pellet was resuspended in 400μL of sorting buffer containing 50μL of MojoSort Streptavidin Nanobeads (BioLegend, 480016) and kept in cold and dark for 15 min. After incubation, an additional 400μL was added to cells, and the tube was placed on EasyEights EasySep Magnet (STEMCELL, 18103) for 7 min. After that supernatant was transferred to a fresh 5mL tube and centrifuged at 600g for 10 min. Pellet was resuspended in 100μL of sorting buffer and 10μL of MojoSort Mouse anti-APC Nanobeads (BioLegend, 480072) was added. The suspension was gently pipetted and incubated for 15 min at 4°C in dark. Afterward, the tube was placed on a magnet for 7 min. Next, the supernatant was saved for analysis, and beads with attached F4/80+ cells were washed with sorting buffer and counted under a light microscope with a Neubauer chamber. Cells were pelleted and frozen in liquid nitrogen for further analysis.

For isolation of extracellular iron-containing aggregates, splenocytes were resuspended in HBSS and then carefully layered over Lymphosep (3:1) in a FACS tube, creating a sharp spleen cell suspension-Lymphosep interphase. Leukocytes were sorted out from the supernatant after density centrifugation at 400g at 20°C for 25 min. The pellet comprising mostly RBCs, granulocytes, and extracellular iron-containing aggregates was then washed again with HBSS to remove Lymphosep. The cell pellet was re-suspended in a sorting buffer and then passed through a magnetized LS separation column (Miltenyi Biotec). The iron-containing superparamagnetic cells/aggregates were eluted from the demagnetized column, washed, and re-suspended in a sorting buffer. To achieve a pure yield of extracellular iron-containing aggregates and remove any trace contaminations from superparamagnetic RPMs or other leukocytes, cells expressing F4/80, B220, Gr-1, CD3, and Ly-6C were sorted out using MojoSort magnetic cell separation system as previously described. The remaining material comprising of mostly aggregates was washed thoroughly and either pelleted and frozen in liquid nitrogen for further analysis (mass spectrometry and iron/heme measurements) or stained with fluorophore-conjugated antibodies for purity verification with flow cytometry.

### Proteomic analyses of splenic aggregates using label-free quantification (LFQ)

#### Sample preparation

Magnetically-isolated aggregates were dissolved in neat trifluoroacetic acid. Protein solutions were neutralized with 10 volumes of 2M Tris base, supplemented with TCEP (8mM) and chloroacetamide (32mM), heated to 95 °C for 5min, diluted with water 1:5, and subjected to overnight enzymatic digestion (0.5μg, Sequencing Grade Modified Trypsin, Promega) at 37 °C. Tryptic peptides were then incubated with Chelex 100 resin (25mg) for 1h at RT, desalted with the use of AttractSPE™ Disks Bio C18 (Affinisep), and concentrated using a SpeedVac concentrator. Prior to LC-MS measurement, the samples were resuspended in 0.1% TFA, 2% acetonitrile in water.

#### LC-MS/MS analysis

Chromatographic separation was performed on an Easy-Spray Acclaim PepMap column 50cm long × 75μm inner diameter (Thermo Fisher Scientific) at 45°C by applying a 90 min acetonitrile gradients in 0.1% aqueous formic acid at a flow rate of 300nl/min. An UltiMate 3000 nano-LC system was coupled to a Q Exactive HF-X mass spectrometer via an easy-spray source (all Thermo Fisher Scientific). The Q Exactive HF-X was operated in data-dependent mode with survey scans acquired at a resolution of 120,000 at m/z 200. Up to 12 of the most abundant isotope patterns with charges 2-5 from the survey scan were selected with an isolation window of 1.3 m/z and fragmented by higher-energy collision dissociation (HCD) with normalized collision energies of 27, while the dynamic exclusion was set to 30s. The maximum ion injection times for the survey scan and the MS/MS scans (acquired with a resolution of 15,000 at m/z 200) were 45 and 96ms, respectively. The ion target value for MS was set to 3e6 and for MS/MS to 1e5, and the minimum AGC target was set to 1e3.

#### Data processing

The data were processed with MaxQuant v. 2.0.3.0 (Cox and Mann, 2008), and the peptides were identified from the MS/MS spectra searched against the reference mouse proteome UP000000589 (Uniprot.org) using the build-in Andromeda search engine. Raw files corresponding to 3 replicate samples obtained from Ag isolates and 3 replicate samples obtained from Y isolates were processed together. Cysteine carbamidomethylation was set as a fixed modification and methionine oxidation, glutamine/asparagine deamidation, and protein N-terminal acetylation were set as variable modifications. For in silico digests of the reference proteome, cleavages of arginine or lysine followed by any amino acid were allowed (trypsin/P), and up to two missed cleavages were allowed. LFQ min. ratio count was set to 1. The FDR was set to 0.01 for peptides, proteins and sites. Match between runs was enabled. Other parameters were used as pre-set in the software. Unique and razor peptides were used for quantification enabling protein grouping (razor peptides are the peptides uniquely assigned to protein groups and not to individual proteins). Data were further analyzed using Perseus version 1.6.10.0 (Tyanova et al., 2016) and Microsoft Office Excel 2016.

#### Data processing and bioinformatics

Intensity values for protein groups were loaded into Perseus v. 1.6.10.0. Standard filtering steps were applied to clean up the dataset: reverse (matched to decoy database), only identified by site, and potential contaminants (from a list of commonly occurring contaminants included in MaxQuant) protein groups were removed. Reporter intensity values were normalized to the tissue weight the aggregates were isolated from and then Log2 transformed. Protein groups with valid values in less than 2 Ag samples were removed. For protein groups with less than 2 valid values in Y, missing values were imputed from a normal distribution (random numbers from the following range were used: downshift = 1.8StdDev, width = 0.4StdDev). One-sided Student T-testing (permutation-based FDR = 0.001, S0 = 1) was performed on the dataset to return 3290 protein groups whose levels were statistically significantly greater in Ag samples compared to Y samples. Annotation enrichment analysis was performed using DAVID (https://david.ncifcrf.gov/) and ShinyGO (http://bioinformatics.sdstate.edu/go/), using FDR=0,05 as a threshold.

### Proteomic analyses of splenic aggregates Tandem Mass Tag (TMT) labeling

#### Sample preparation

Magnetically-isolated aggregates were dissolved in neat trifluoroacetic acid. Protein solutions were neutralized with 10 volumes of 2M Tris base, supplemented with TCEP (8mM) and chloroacetamide (32mM), heated to 95 °C for 5min, diluted with water 1:5, and subjected to overnight enzymatic digestion (0.5μg, Sequencing Grade Modified Trypsin, Promega) at 37 °C. Tryptic peptides were then incubated with Chelex 100 resin (25mg) for 1h at RT, desalted with the use of AttractSPE™ Disks Bio C18 (Affinisep), TMT-labeled on the solid support (Myers et al., 2019), compiled into a single TMT sample and concentrated using a SpeedVac concentrator. Prior to LC-MS measurement, the samples were resuspended in 0.1% TFA, 2% acetonitrile in water.

#### LC-MS/MS analysis

Chromatographic separation was performed on an Easy-Spray Acclaim PepMap column 50cm long × 75μm inner diameter (Thermo Fisher Scientific) at 45°C by applying a 120 min acetonitrile gradients in 0.1% aqueous formic acid at a flow rate of 300nl/min. An UltiMate 3000 nano-LC system was coupled to a Q Exactive HF-X mass spectrometer via an easy-spray source (all Thermo Fisher Scientific). Three samples injections were performed. The Q Exactive HF-X was operated in data-dependent mode with survey scans acquired at a resolution of 60,000 at m/z 200. Up to 15 of the most abundant isotope patterns with charges 2-5 from the survey scan were selected with an isolation window of 0.7 m/z and fragmented by higher-energy collision dissociation (HCD) with normalized collision energies of 32, while the dynamic exclusion was set to 35s. The maximum ion injection times for the survey scan and the MS/MS scans (acquired with a resolution of 45,000 at m/z 200) were 50 and 96ms, respectively. The ion target value for MS was set to 3e6 and for MS/MS to 1e5, and the minimum AGC target was set to 1e3.

#### Data processing

The data were processed with MaxQuant v. 1.6.17.0 (Cox and Mann, 2008), and the peptides were identified from the MS/MS spectra searched against the reference mouse proteome UP000000589 (Uniprot.org) using the build-in Andromeda search engine. Raw files corresponding to 3 replicate injections of the combined TMT sample were processed together as a single experiment/single fraction. Cysteine carbamidomethylation was set as a fixed modification and methionine oxidation, glutamine/asparagine deamidation, and protein N-terminal acetylation were set as variable modifications. For in silico digests of the reference proteome, cleavages of arginine or lysine followed by any amino acid were allowed (trypsin/P), and up to two missed cleavages were allowed. Reporter ion MS2 quantification was performed with the min. reporter PIF was set to 0.75. The FDR was set to 0.01 for peptides, proteins and sites. Match between runs was enabled and second peptides function was disabled. Other parameters were used as pre-set in the software. Unique and razor peptides were used for quantification enabling protein grouping (razor peptides are the peptides uniquely assigned to protein groups and not to individual proteins). Data were further analyzed using Perseus version 1.6.10.0 (Tyanova et al., 2016) and Microsoft Office Excel 2016.

#### Data processing and bioinformatics

Reporter intensity corrected values for protein groups were loaded into Perseus v. 1.6.10.0. Standard filtering steps were applied to clean up the dataset: reverse (matched to decoy database), only identified by site, and potential contaminants (from a list of commonly occurring contaminants included in MaxQuant) protein groups were removed. Reporter intensity values were Log2 transformed and normalized by median subtraction within TMT channels. 942 Protein groups with the complete set of valid values were kept. Student T-testing (permutation-based FDR = 0.05, S0 = 0.1) was performed on the dataset to return 70 protein groups, which levels were statistically significantly changed in Ag vs IR samples. Annotation enrichment analysis was performed using DAVID (https://david.ncifcrf.gov/) and ShinyGO (http://bioinformatics.sdstate.edu/go/), using FDR=0,05 as a threshold.

### Measurement of cellular iron levels

Determination of cellular total iron levels in magnetically-sorted RPMs was carried out using the Iron assay Kit (Sigma-Aldrich, MAK025) according to the manufacturer’s instructions, and as shown previously (Folgueras et al., 2018). Iron concentrations (ng/μL) were calculated from the standard curve and normalized to the number of cells for each sample.

### RBCs *in vivo* lifespan

EZ-Link Sulfo-NHS Biotin (Thermo Scientific, 21217) was dissolved in sterile PBS to a final concentration of 1mg per 100μL and filtered through a 0,1μm filter (Millipore, SLVV033RS). A day before the first blood collection, 100μL of the sterile solution was injected intravenously into mice. On days 0, 4, 11, 18, and 25 approximately 10 μL of whole blood was collected from the tail vein with heparinized capillary to a tube containing HBSS. RBCs were centrifuged at 400g for 5 min at 4°C. Each sample was resuspended in 250μL of HBSS containing 5% normal rat serum (Thermo Scientific). Then 2μL of fluorescently labeled anti-TER-119 and streptavidin was added to the suspension. Fluorescent streptavidin was omitted for FMO samples in each group. After incubation at 4°C for 30 min, samples were centrifuged and resuspended with HBSS. The percentage of biotinylated erythrocytes was determined by flow cytometry.

### Preparation of stressed erythrocytes for erythrophagocytosis assays

Preparation and staining of stressed RBCs (sRBCs) were performed as described before (Theurl et al., 2016), with some modifications. **Preparation of RBCs:** Mice were sacrificed and whole blood was aseptically collected *via* cardiac puncture to CPDA-1 solution (Sigma-Aldrich, C4431). The final concentration of CPDA-1 was 10%. Whole blood obtained from mice was pooled and then centrifuged at 400g for 15 min at 4°C. Plasma was collected and filtered through a 0.1μm filter and stored at 4°C. RBCs were resuspended in HBSS and leukoreduced using Lymphosep (Biowest, L0560-500). Cells were washed with HBSS and then heated for 30 min at 48°C while continuously shaking, generating sRBC. **Staining of sRBCs:** 1 × 10^10^ RBC were resuspended in 1 ml diluent C, mixed with 1 ml diluent C containing 4μM PKH-67 (Sigma-Aldrich, MIDI67-1KT) and incubated in dark for 5 min in 37°C, the reaction was stopped by adding 10mL HBSS containing 2% FCS and 0.5% BSA. Staining was followed by two washing steps with HBSS. For *in vitro* and *ex vivo* erythrophagocytosis assay cells were resuspended in RPMI-1640 and counted.

For *in vivo* approach, RBCs were resuspended to 50% hematocrit in previously collected and filtered plasma.

### *In vitro* erythrophagocytosis

Stained and counted sRBCs were added to iRPMs on 12-well plates in 10 fold excess for 1.5h in 37°C, 5% CO_2_ on the 8^th^ day after seeding. After that cells were extensively washed with cold PBS to discard not engulfed sRBCs. Next, cells were detached with Accutase (BioLegend, 423201), transferred to a round bottom tube, washed with HBSS, and centrifuged at 600g for 5 min. Cells were labeled with antibodies and analyzed by flow cytometry.

### *In vivo* erythrophagocytosis

Mice were injected into the tail vein with 100μL of RBCs resuspended in plasma to 50% hematocrit. Mice were maintained for 1.5h in cages with constant access to water and food. After that time animals were sacrificed for organ isolation.

### *Ex vivo* phagocytosis and erythrophagocytosis

10×10^6^ splenocytes were resuspended in a 5mL round bottom tube in 200μL of warm complete RPMI-1640. Fluorescent sRBCs or fluorescent Zymosan A particles (Invitrogen, Z23373) were added to cells at ratio 10:1 (sRBCs/Zymosan: Cells) for 1.5h at 37°C, 5% CO_2_. Afterward, cells were centrifuged at 600g for 5 min. Excess of sRBCs was lysed using 1X RBCs lysis buffer, cells were washed with HBSS and centrifuged. Next, cells were labeled with fluorescent antibodies and analyzed by flow cytometry.

### Heme content analysis

The whole spleen was weighted, quickly dissected and gently mashed through a 100μm strainer in the presence of 3mL HBSS. After that suspension was centrifuged at 400g for 10 min at 4°C. A splenocyte pellet was used for different purposes and 1mL of heme-containing supernatant was transferred to a 1,5mL tube and centrifuged at 1000g for 10 min at 4°C.

Splenic aggregates were isolated as previously described and were suspended in ddH_2_O water. Heme concentration was measured using Heme Assay Kit (Sigma-Aldrich, MAK316) according to manufacturer instructions. Absorbance was measured at 400nm. The amount of heme was calculated against Heme Calibrator and normalized to the initial weight of fresh spleens.

### Transferrin saturation and tissue iron measurements

Serum iron and unsaturated iron-binding capacity were measured with SFBC (Biolabo, 80008) and UIBC (Biolabo, 97408) kits according to manufacturer protocols. Transferrin saturation was calculated using the formula SFBC/(SFBC+UIBC)x100. For measurement of tissue non-heme iron content, the bathophenanthroline method was applied and calculations were made against dry weight tissue, as described previously (Torrance and Bothwell, 1968).

### Erythropoietin (EPO) measurement with ELISA

The plasma levels of erythropoietin were measured by Mouse Erythropoietin/EPO Quantikine ELISA Kit (R&D Systems, MEP00B) according to the manufacturer’s instructions. The optical density was measured on a microplate reader at a wavelength of 450nm with wavelength correction set to 570nm.

### RNA isolation

RNA from sorted cells was isolated from TRIzol LS Reagent (Invitrogen, 10296028) using Direct-zol RNA Microprep Kit (Zymo Research, R2062) according to the manufacturer’s instructions. RNA from tissues was isolated from TRIzol Reagent (Invitrogen, 15596018) following the guidelines of the manufacturer protocol.

### Reverse transcription and qRT-PCR

cDNA was synthesized with RevertAid H Minus Reverse Transcriptase (Thermo Scientific, EP0452) according to manufacturer guidelines. Real-time PCR was performed by using SG qPCR Master Mix (EURx, E0401) and HAMP gene primers (Forward 5’-ATACCAATGCAGAAGAGAAGG-3’, Reverse 5’-AACAGATACCACACTGGGAA-3’) as described in manufacturer protocol. qRT-PCR was run on LightCycler 96 System (Roche).

### Histological and histochemical analysis

Following fixation in 10% formalin for 24 h, spleens were stored in 70% ethanol before further preparation. The tissue was embedded in paraffin and 7-μm cross-sections were cut with a microtome (Reichert-Jung, Germany). The sections were stained with hematoxylin and eosin. Slides were examined by light microscopy (Olympus, type CH2). Non-heme iron staining of spleen samples was analyzed using Accustain Iron Deposition Kit (Sigma). Sections were prepared as described above. After mounting on glass slides, sections were deparaffinized, incubated with a working solution containing Perls’ Prussian Blue for 30 min, counterstained with pararosaniline solution for 2 min and analyzed under standard light microscopy (Olympus CH2).

### Transmission electron microscopy (TEM) of the spleen

Fresh samples of the spleen, about 3 square mm, were fixed in 2.5% glutaraldehyde for 24 h at 4°C, then washed in PBS and postfixed with 1 % osmium tetroxide for 1 h. After washing with water they were incubated with 1% aqueous uranyl acetate for 12 h at 4°C. Next, samples were dehydrated at room temperature with increasing concentrations of ethanol, infiltrated with epoxy resin (Sigma Aldrich, St. Louis, MO, USA, #45-359-1EA-F) and subjected for polymerization for 48 h at 60°C. Polymerized resin blocks were trimmed with a tissue processor (Leica EM TP), cut with an ultramicrotome (EM UC7, Leica) for ultrathin sections (65 nm thick), and collected on nickel grids, mesh 200 (Agar Scientific, G2200N). Specimen grids were examined with a transmission electron microscope Tecnai T12 BioTwin (FEI, Hillsboro, OR, USA) equipped with a 16 megapixel TemCam-F416 (R) camera (TVIPS GmbH) at in-house Microscopy and Cytometry Facility.

### Transcriptome analysis by RNA-seq

To prepare libraries from FACS-sorted RPMs (at least 100 000 cells/sample), we used the previously described Smart-seq2 protocol (Picelli et al., 2013), suitable for low-input total mRNA sequencing. The quality of RNA and material during the preparation of libraries was checked by Bioanalyzer. The samples were sequenced on NextSeq500 (Illumina) with 75 bp single-end reads, with ~50 million reads/sample. RNAseq was performed at GeneCore at EMBL (Heidelberg, Germany). The quality of the reads was assessed with FastQC software [https://www.bioinformatics.babraham.ac.uk/projects/fastqc/]. Reads were mapped to the Mus musculus genome assembly GRCm38(mm10) with HISAT2 software [http://daehwankimlab.github.io/hisat2/] on default parameters. Then, the mapped reads were counted into Ensembl annotation intervals using HTSeq-count software [https://www.ncbi.nlm.nih.gov/labs/pmc/articles/PMC4287950/]. Differentially expressed genes were estimated using DESeq2 software [https://www.ncbi.nlm.nih.gov/labs/pmc/articles/PMC4302049/] with default parameters. Genes with p-adjusted < 0.05 were regarded as differentially expressed and included in further analysis. Functional analysis was conducted using ClusterProfiler [https://doi.org/10.1016/j.xinn.2021.100141].

### Statistical analysis

Female mice of the same age were randomly attributed to experimental groups (diets or NAC administration). Mouse-derived samples were collected randomly within groups and often assessed/measured randomly, and groups were harvested in a different order in individual experiments. The investigators were not blinded during experiments and assessments. Sample size (typically 4-7 mice/group or independent biological cell-based experiments) was determined based on power analysis, prior experience of performing similar experiments, and previously published papers in the field of RPM biology (Akilesh et al., 2019; Lu et al., 2020; Ma et al., 2021; Okreglicka et al., 2021). Statistical analysis was performed with GraphPad Prism (GraphPad software, Version 9). Data are represented as mean ± SEM, unless otherwise specified. ROUT method was applied (in rare cases) to identify and remove outliers. For all experiments, α=0.05. When two groups were compared two-tailed unpaired Welch’s t-test was applied, whereas for multiple comparisons, the One-Way Analysis of Variance (ANOVA) test was performed. For ANOVA, Dunnett’s Multiple Comparison test was used for experiments comparing multiple experimental groups to a single control, while post-hoc Tukey’s test was used to compare multiple experimental groups. The number of mice/samples per group or the number of independent cell-based experiments are shown in the figures. Results were considered as significant for p< 0.05, ** - p (* - p< 0.05, ** - p<0.01, *** -p<0.001, ****- p<0.0001).

## Supporting information

Table S1. Proteomics LFQ analysis.

Table S2. Proteomics TMT analysis.

## Data availability

RNA sequencing data are deposited in the GEO repository (under accession no: GSE199879). The following secure token has been created to allow review of record while it remains in private status: izgfeykoxjyzlyt. Mass spectrometry proteomics data were deposited to the ProteomeXchange Consortium via the PRIDE partner repository with the dataset identifier PXD032900. These data can be accessed via the Reviewer account: Username; Password: Z4AfdopG.

## Author contribution

Conceptualization: PS, PKM, WP, KMS; Formal analysis: PS, PKM, GZ, OK, MMa, AS, MM, ML, RS, WP, KMS; Funding acquisition: MM, ML, WP, KMS, Investigation: PS, PKM, GZ, MN, KKC, SH, MMa, AS, ML, KMS; Methodology; PS, KPM, GZ, MC, OK, SH, MMa, AS, ML, RS, KMS; Project administration: WP, KMS; Resources: WP, KMS; Supervision: WP, KMS; Validation: MM, ML, WP, KMS; Visualization: PS, PKM, GZ, OK, MMa, ML, KMS; Writing – original draft: PS, PKM, KMS; Writing – review & editing: WP, KMS.

## Acknowledgments

We thank the GeneCore team (EMBL, Heidelberg) for performing RNA sequencing. We thank Tara Arvedson (Amgen Inc., USA) for the anti-ferroportin antibody. Many thanks to Agnieszka Popielska and the staff of the Experimental Medicine Centre (Bialystok, Poland) for their technical support. Proteomic measurements were performed at the Proteomics Core Facility, IMol Polish Academy of Sciences utilizing the equipment funded by the ‘Regenerative Mechanisms for Health’ project MAB/2017/2 within the International Research Agendas program of the Foundation for Polish Science, co-financed by the European Union under the European Regional Development Fund. We thank Dr. Dorota Stadnik for the preparation and measurement of the proteomic samples and Dawid Hatala for his assistance in histological analyses. We acknowledge the help of Raffaella Gozzellino in the initial phase of project conceptualization. WP and KMS acknowledge internal IIMCB funding for inter-lab projects. WP acknowledges funding from Norwegian Financial Mechanism 2014-2021 and operated by the Polish National Science Center under the project contract no UMO-2019/34/H/NZ3/00691 and European Molecular Biology Organization (EMBO Installation Grant No. 3916). KMS acknowledges funding from the National Science Centre Sonata Bis grant (UMO-2020/38/E/NZ4/00511). The model figure was prepared using BioRender.com.

## Competing interests

The authors declare no conflict of interest.

## Notes

### Competing Interest Statement

The authors have declared no competing interest.

### Summary of Updates

This is a significantly revised version of our manuscript updated with the data generated over last 2 months.

