## Supplementary Figures for "Impaired iron recycling from erythrocytes is an early hallmark of aging"

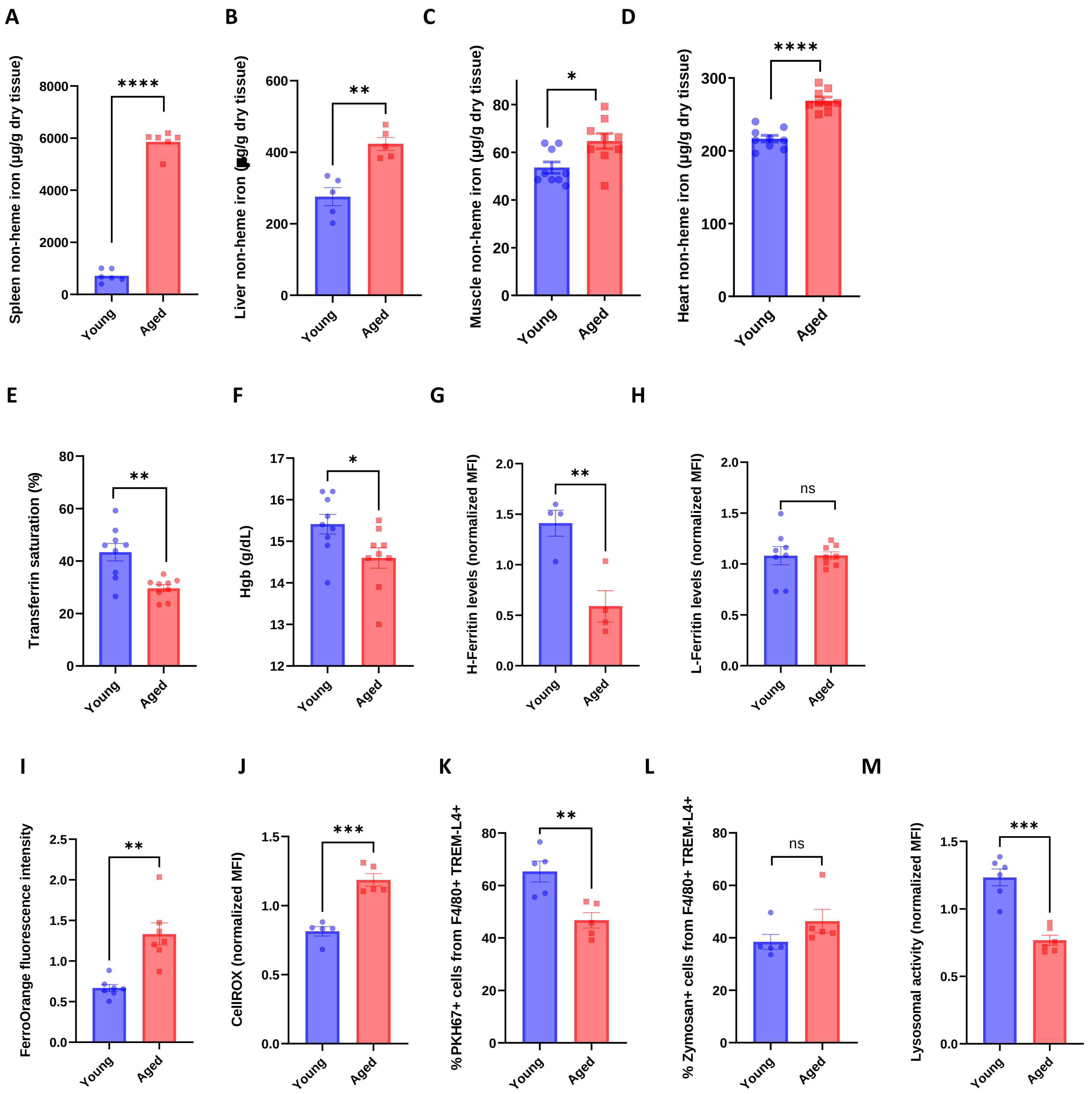


Figure S1. **RPMs of aged mice show increased labile iron levels, oxidative stress and diminished iron-recycling functions.**

(A-D) Spleen, liver, muscle and heart non-heme iron content was determined in young and aged mice.

(E) Plasma transferrin saturation and (F) blood hemoglobin (Hgb) concentration were determined in young and aged mice.

(G) In young and aged RPMs, H-Ferritin and (H) L-Ferritin were quantified with flow cytometric analysis.

(I) Cytosolic ferrous iron ($Fe2+$) and (J) ROS levels in young and aged RPMs were quantified using the FerroOrange and CellROX Deep Red probes, respectively, and flow cytometry.

(K) The phagocytosis rates of PKH67-labeled temperature-stressed RBCs and (L) zymosan A fluorescent particles by young and aged RPMs were analyzed *ex vivo* by flow cytometry. The percent of cargo-positive RPMs is indicated.

(M) Lysosomal activity in young and aged RPMs was determined using a dedicated fluorescent probe and flow cytometry.

Each dot represents one mouse. Data are represented as mean ± SEM. Welch's unpaired t-test determined statistical significance between the two groups. *p<=0.05, **p<=0.01, ***p<=0.001 and ****p<=0.0001

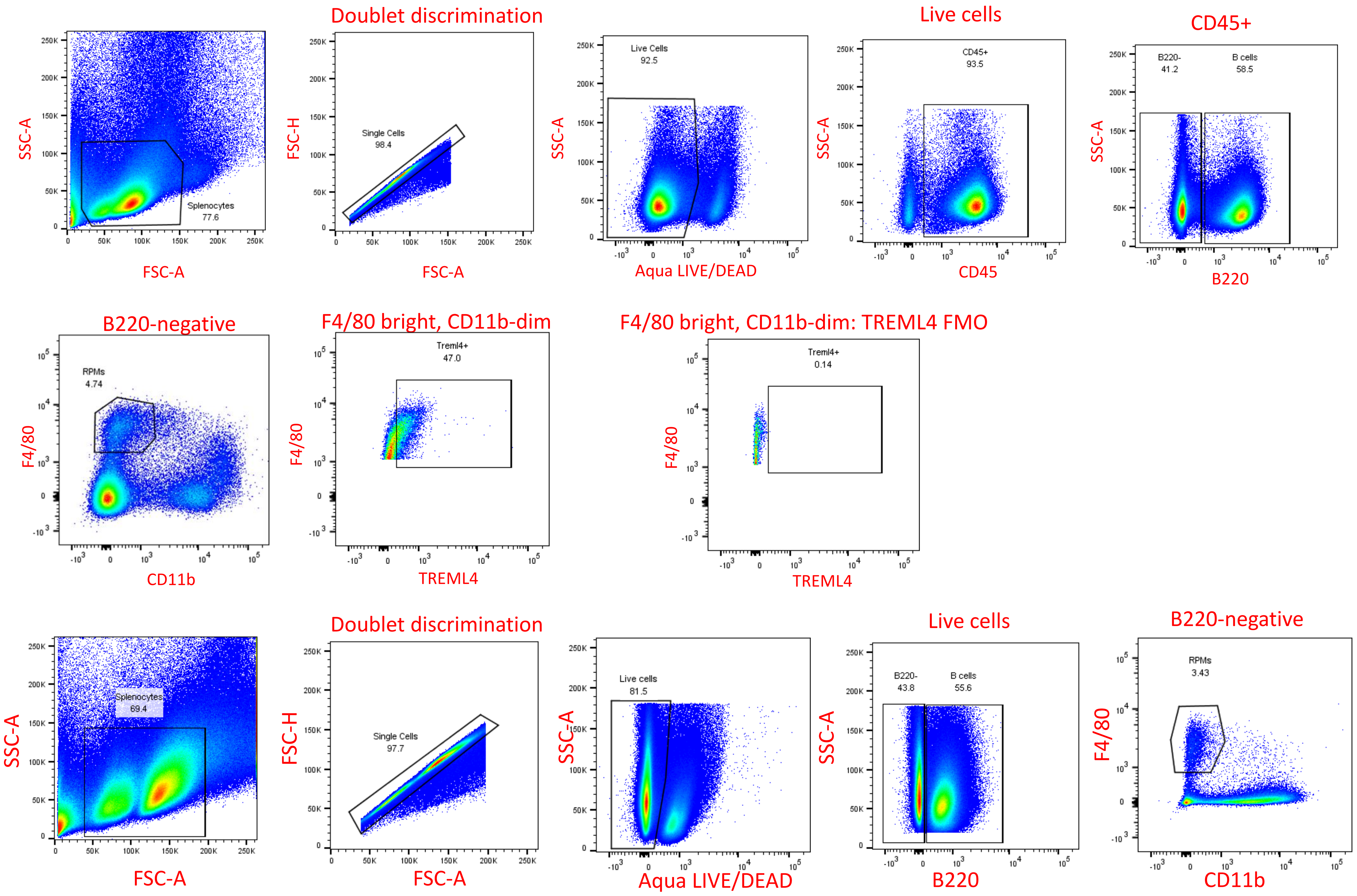


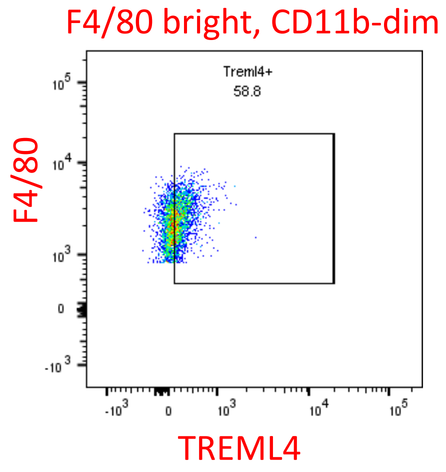


Figure S2. **Gating strategy for RPMs.**

(A) Non-permeabilized cells are shown, including an FMO control .

(B) Permeabilized cells are shown

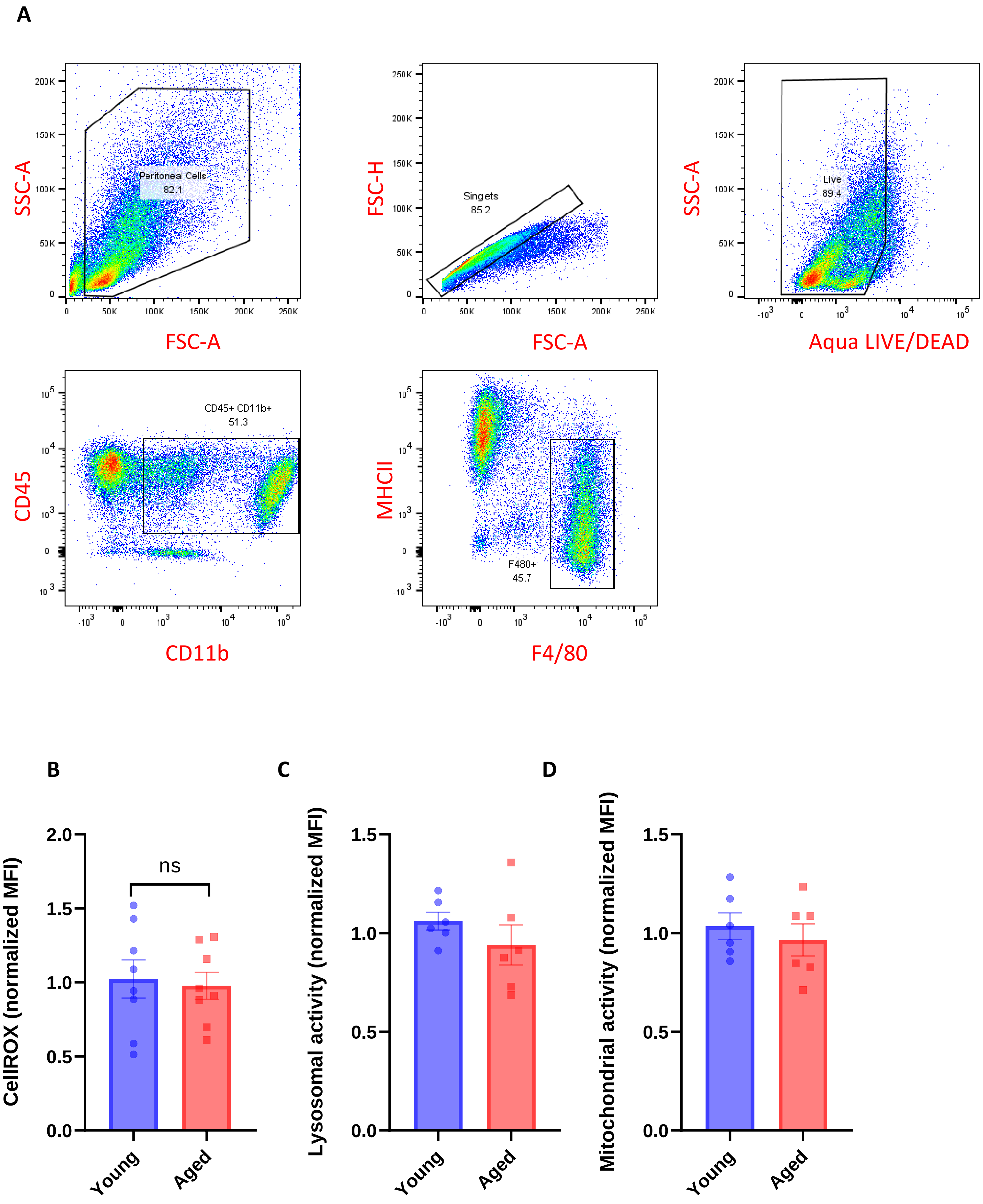


Figure S3. **Peritoneal macrophages in aged mice do not show functional impairments.**

(A) Gating strategy for peritoneal macrophages.

(B) ROS levels in young and aged peritoneal macrophages were quantified using CellRox Deep Red with flow cytometry.

(C) A dedicated fluorescent probe and flow cytometry determined lysosomal activity in young and aged peritoneal macrophages.

(D) The mitochondrial membrane potential in young and aged peritoneal macrophages was measured by staining with TMRE and flow cytometry analysis.

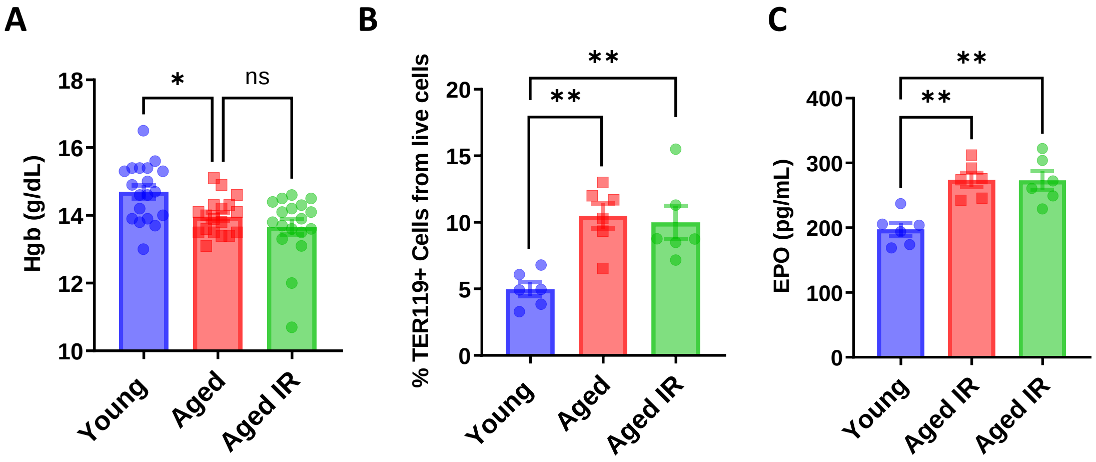


Figure S4. **Mild anemia of aged mice is not rescued by the iron-reduced diet**.
(A) Blood hemoglobin (Hgb) concentration was determined in young, aged, and aged IR mice.
(B) Shown is the percentage of erythroid cells (TER119+) present in the spleen and (C) EPO concentration in the plasma of young, aged, and aged IR mice.

Each dot represents one mouse. Data are represented as mean ± SEM. Statistical significance among the three groups was determined by the One-Way ANOVA test with Tukey's Multiple Comparison test. *p<=0.05, **p<=0.01.

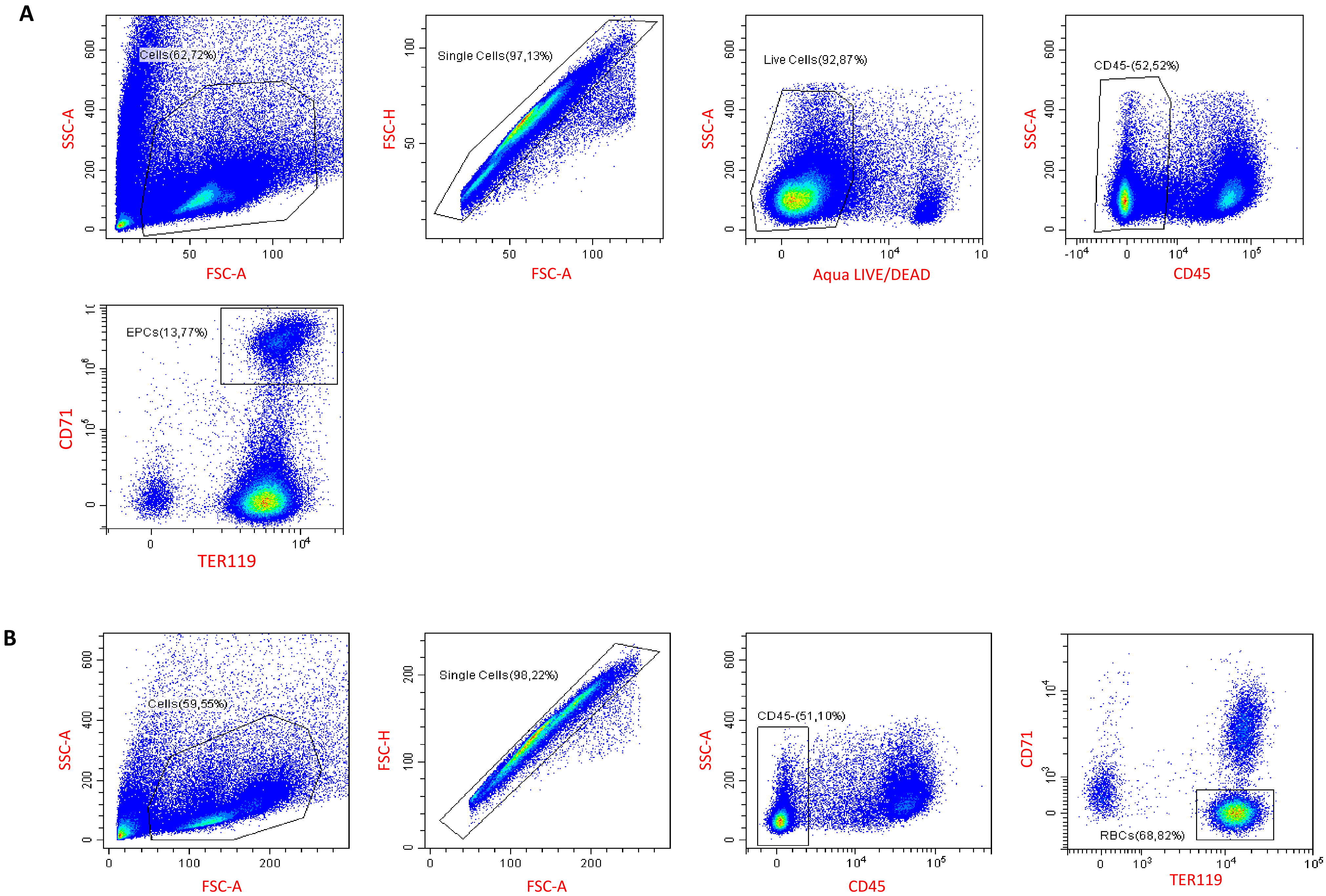


Figure S5. **Gating strategy for erythroid progenitor cells (A) and splenic RBCs (B)**

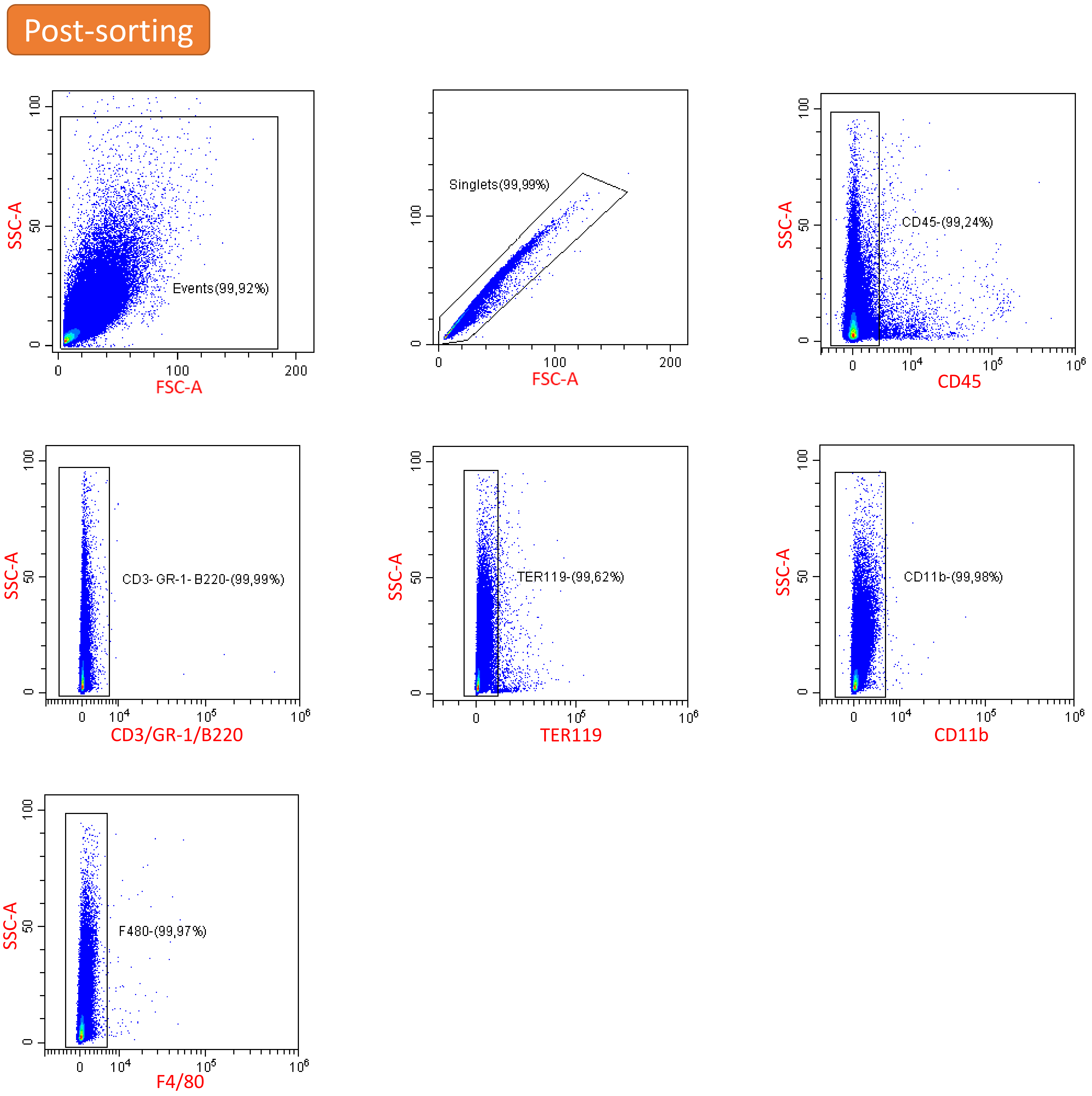


Figure S6. **Post-sorting purity of extracellular aggregates isolated from the aged spleen**. Shown is the flow cytometry analysis of the isolated aggregates that are in >99% negative for the splenocyte cell markers.

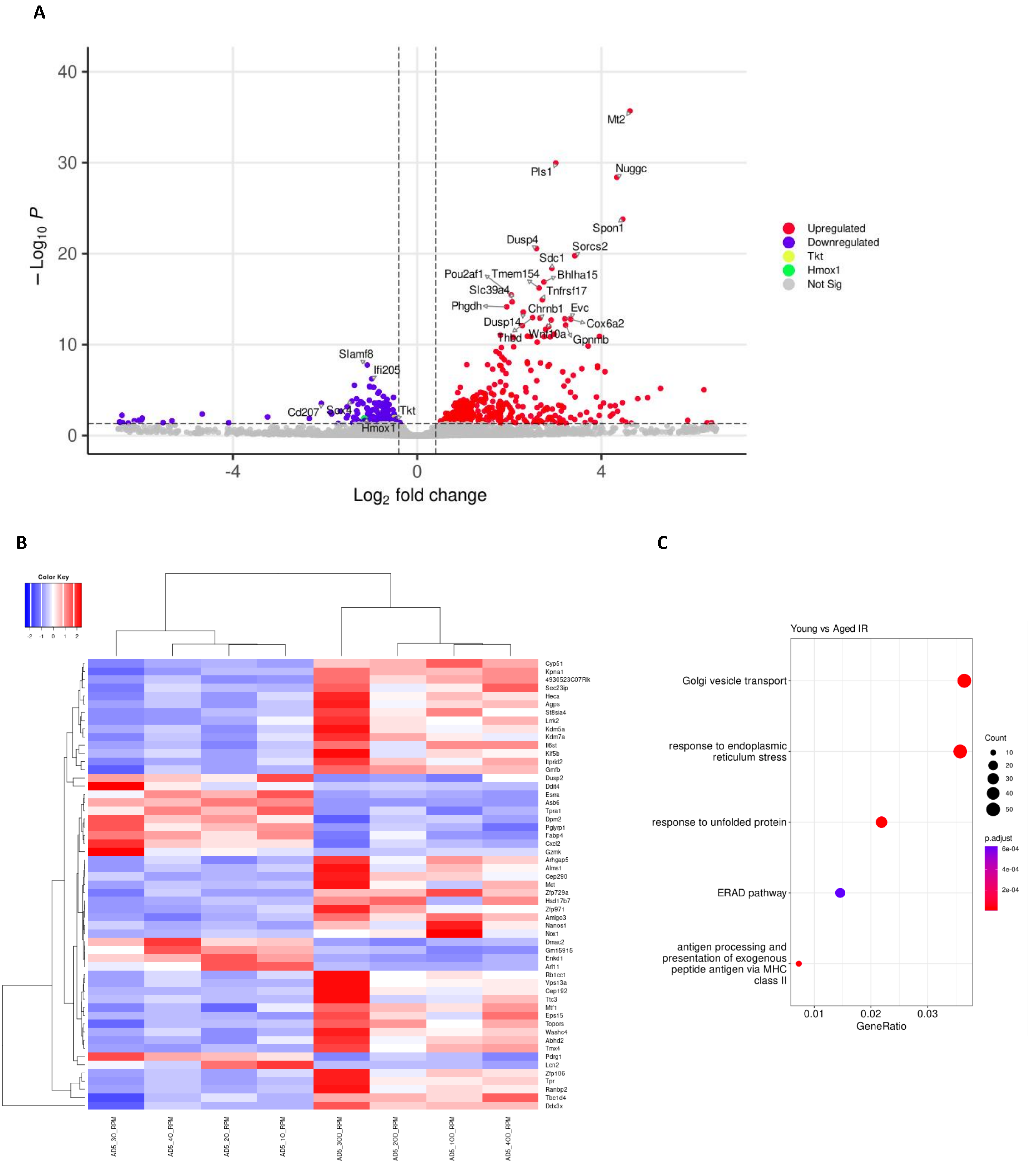


Figure S7. **Supplementary data related to RNA-seq analysis of RPMs derived from young, aged and aged IR mice** (related to Figure 6).

(A) Volcano plot of differentially regulated genes in aged *versus* young RPMs, with top hits indicated.

(B) Heat map representing 54 differentially regulated genes between aged and aged IR RPMs.

(C) Enriched functional categories among differentially regulated genes in FACS-sorted RPMs derived from young *versus* aged IR mice
